# Characterization of divalent cation interactions with AASTY native nanodiscs

**DOI:** 10.1101/2021.10.07.463511

**Authors:** Milena Timcenko, Anton A. A. Autzen, Henriette E. Autzen

## Abstract

Amphiphilic copolymers show promise in extracting membrane proteins directly from lipid bilayers into ‘native nanodiscs’. However, many such copolymers are polyanionic and sensitive to divalent cations, limiting their applicability. We characterize the Ca^2+^ and Mg^2+^ sensitivity of poly(acrylic acid-co-styrene) (AASTY) copolymers with analytical UV and fluorescent size exclusion chromatography, enabling us to separate signals from nanodiscs, copolymers, and soluble aggregates. We find that divalent cations promote aggregation and precipitation of both free and lipid bound copolymers. We see that excess, free copolymer acts as a ‘cation sink’ that protects nanodiscs from Ca^2+^ induced aggregation. Removal of the free copolymer through dialysis induces aggregation that can be mitigated by KCl. Finally, we find that the nanodisc size is dynamic and dependent on lipid concentration. Our results offer insight to nanodisc behaviour, and can help guide experimental design, aimed at mitigating the shortcomings inherent in negatively charged nanodisc forming copolymers.

## Introduction

Membrane proteins are involved in numerous cellular functions and pathways and are important therapeutic targets for a variety of diseases. Characterization of integral membrane proteins by biochemical and biophysical methods often requires their isolation from their native lipid environment. This remains a major bottleneck as initial extraction of membrane proteins from the membrane is typically carried out through solubilization with detergents. However, detergent solubilization can be too harsh for membrane proteins or complexes as the detergent strips the majority of endogenous lipids and cofactors from the proteins.^1–3^ Several approaches for re-lipidating the detergent solubilized membrane protein and reconstituting it into a bilayer mimicking environment have been developed, including membrane scaffold protein (MSP) nanodiscs, Saposin-lipoprotein (Salipro) nanoparticles and peptidiscs.^4–7^ In recent years, amphiphilic copolymers have emerged as an attractive alternative to detergents, as they enable direct solubilization of membrane proteins encapsulated by a patch of the surrounding lipid bilayer from the membrane into so-called native nanodiscs.^2,7^ With the native nanodisc approach, the membrane protein remains lipidated, and endogenous lipids and cofactors are preserved around it. Styrene-maleic acid (SMA) copolymers (Figure 1A) were the first to be used for generation of native nanodiscs, and their use remains widespread.^8,9^ While the native nanodisc approach shows great promise, there are significant shortcomings with the currently used copolymers; These include their negative charge, and heterogeneity in terms of molecular weight and monomer sequence, which altogether limit the applicability of the method to a wider range of protein targets.^10^ Alternative copolymers which try to ameliorate these challenges have been published, including poly(diisobutylene-co-maleic acid) (DIBMA),^11^ polymethacrylate (PMA),^12^ stilbene–maleic anhydride (STMA)^13^ and various SMA derivative copolymers^14–17^, as well as a non-charged inulin derivative.^18^ The current developments in native nanodiscs have recently been reviewed by e.g. Brown *et al*.^2^ and Esmaili *et al*.^9^

**Figure 1:**
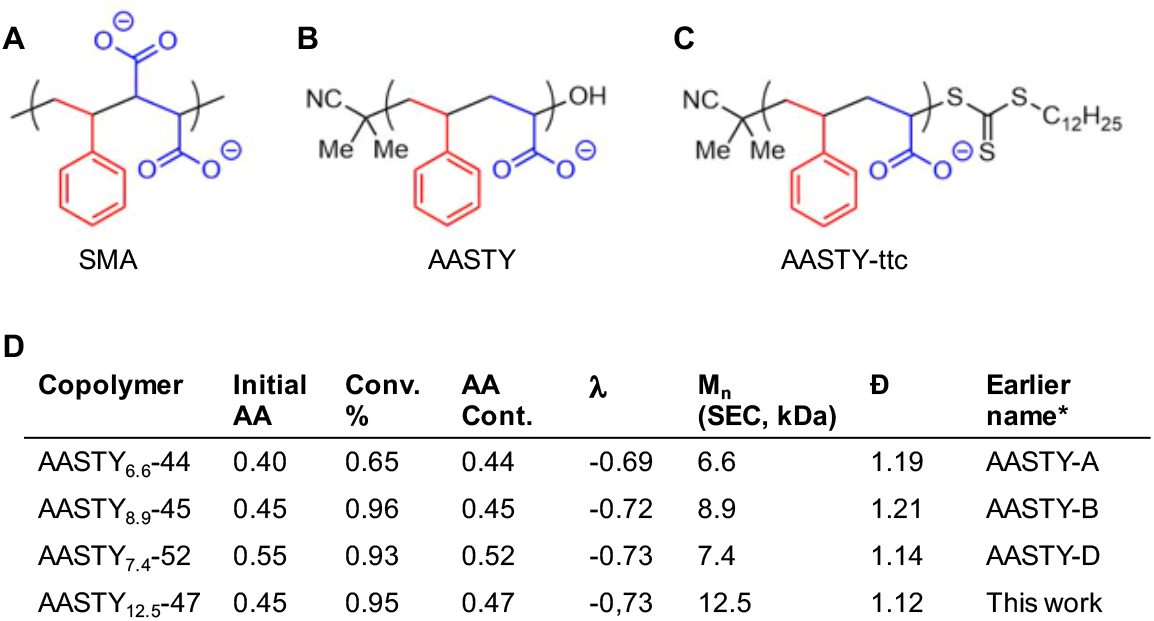
Overview of copolymers used in this study. Chemical structures of **A)** SMA, **B)** AASTY, and **C** AASTY-ttc with the ttc end group intact, which absorbs strongly at 310 nm. This end group comes from the RAFT chain transfer agent used during synthesis and is removed in the regular AASTY. **D)** AASTY copolymer characteristics. Abbreviations and symbols used: Acrylic acid, AA; conversion, Conv.; acrylic acid content, AA Cont.; Chemical correlation parameter, λ (λ = 0 is a perfectly random copolymer, λ = –1 is perfectly alternating); Number average molecular weight, M_n_; dispersity, D= M_w_/M_n_, where M_w_ is the weight average molecular weight. *The name used in Smith *et al*.^19^

We previously showed that poly(acrylic acid-co-styrene) (AASTY) copolymers (Figure 1B) are effective at solubilizing small, unilameller vesicles (SUVs) as well as extracting a model membrane protein from HEK293 cells.^19^ AASTY can be synthesized by reversible addition-fragmentation chain-transfer (RAFT) polymerizations which allows control of copolymer length and dispersity of size, compared with free-radical polymerization.^19–21^

Due to the often polyanionic nature of many copolymers including AASTY, one challenge is a sensitivity towards divalent cations such as calcium (Ca^2+^) and magnesium (Mg^2+^). The sensitivity is caused by interactions between the cations and the negatively charged carboxylate. It has also been shown that monovalent cations increase yields in purification of membrane proteins in nanodiscs by affinity purification. ^19,22,23^ This suggests that cation interactions with the carboxylate can either be detrimental, or beneficial for nanodisc stability, and that salt composition can be optimized for a given sample. Both Ca^2+^ and Mg^2+^ ions are important for a multitude of cellular processes and are required by many protein for function or integrity.^24,25^ Although, the sensitivity of polyanionic copolymers towards divalent cations is well-known, systematic characterization of the effects of divalent cations on amphiphilic copolymers, nanodiscs and the formation of nanodiscs, is limited. The resistance of amphiphilic co-polymers towards Ca^2+^ has typically been characterized through visual inspection, assessment of turbidity, dynamic light scattering (DLS).^14,17,26^ These are low-resolution techniques, where subtle changes in the solution might not be evident, although these changes could affect the properties of the copolymer. Furthermore, as these techniques record properties of an ensemble of species in solution, the signal from nanodiscs and free copolymer remaining in the solution cannot be separated from one another. Recently, Ravula *et al*. used NMR to characterize the effects of Ca^2+^ and Mg^2+^ ions finding rouleaux stacking of nanodiscs with divalent cations. ^27^ This finding further highlights the benefit of in-depth characterization of these systems.

An often cited disadvantage of styrene based copolymers such as AASTY and SMA is their strong absorption of UV light overlapping with the absorption of proteins at 280 nm, which is commonly used for determination of protein concentration and purity.^11^ However, in this study we use this property to our advantage: Using a combination of a photodiode-array (PDA) detector, which records absorption at 190-800 nm, and fluorescent size exclusion chromatography (FSEC), we are able to separate different species not only by size, but also their spectral properties in a straightforward and robust setup. By recording separate signals from fluorescent lipids in nanodiscs and the styrene rich copolymer AASTY, we can examine the effects of divalent cations on both the free copolymer and assembled nanodiscs. Along with UV and FSEC we utilize a colorimetric assay to examine the interactions between the AASTY copolymers and divalent cations. We find that copolymer resistance towards Ca^2+^ and Mg^2+^ ions depends greatly on the copolymer concentration assayed, where higher concentrations are more tolerant. This relates to the binding capacity of the copolymers for the divalent cations, where higher copolymer concentrations result in lower free Ca^2+^ ion concentration compared with a lower copolymer concentration. Binding of divalent cations thereby appear to yield protection against precipitation to some extent. Likewise, including salt in the buffer also yields protection. We further observe that small changes in the AASTY copolymer composition can have a great effect on their behaviour, where higher acrylic acid content results in higher divalent cation tolerance, though the size of the copolymer also play a role. We find that interactions between AASTY copolymers and divalent cations are highly dependent on the relative concentrations of divalent cations, the copolymer in solution, the copolymer contained in nanodiscs and the ionic strength of the solution. As such, defining a one-fit-all concentration threshold for Ca^2+^ or Mg^2+^ necessary to precipitate the copolymer from solution is not possible.

## Experimental

### Synthesis and characterization of AASTY copolymers

The copolymers (Figure 1) were the same material as used in Smith *et al*.,^19^ with the exception of AASTY_12.5_-47. The synthesis of AASTY_12.5_-47 was analogous to the other copolymers and their characteristics are summarized in Figure 1D. In short, a schlenck flask was charged with azobisisobutyrunitrile (104 mg, 0.634 mmol), the RAFT agent 2-cyano-2-propyl dodecyl trithiocarbonate (1.10 g, 3.17 mmol), and distilled acrylic acid (13.0 g, 190 mmol) and styrene (26.6 g, 232 mmol). The reaction mixture was subjected to 4 freezepump-thaw cycles to remove oxygen, and backfilled with nitrogen on a schlenk line. The reaction was heated to 70° C for 9 hours, reaching 95% monomer conversion, resulting in a yellow solid. The solid was dissolved in diethyl ether, and precipitated into hexane, followed by drying in vacuo, resulting in a yellow crisp solid. The dodecyl trithiocarbonate end group (ttc) was removed by dissolving copolymer (5 g) in a 1:3 mixture of water:ethanol (40 mL) and 30% H_2_O_2_ (1.8 mL), and the mixture was incubated at 70°C overnight, resulting in a colorless solution. The product was precipitated by addition to water, and the product was collected by centrifugation. For conversion to a partial sodium salt, the white solid was mixed with water, and NaOH (1 M) was added until the pH is stable at 7.3. The opaque mixture was filtered and lyophilized to yield the partial sodium salt of AASTY. The characterization of AASTY_12.5_-47 was analogous to the copolymers included in the previous study.^19^ Values for all four polymers are summarized in Figure 1D. In short, the monomer conversion was calculated by ^1^H-NMR (Supp. Figure S1) measured on a Bruker AVANCE 400 MHz system. The number average molecular weight, M_n_ and dispersity, D= M_n_/M_w_, were measured using a Dionex Ultimate 3000 instrument outfitted with a Dawn Heleos II Multi Angle Light Scattering detector, and a Optilab rEX refractive index detector. The column was a Superose 6 Increase column (10/300, Cytiva). Data were analyzed using Astra 7.0 software, using a dndc of 0.170 mL/g..

### Determination of free Ca^2+^ concentrations

We previously showed that the charge state of the AASTY copolymer depend on the pH of the solution.^19^ As many membrane proteins function at a pH of 7.4, we decided to conduct our analysis at this pH value. As such, all experiments with the polymers were carried out at pH 7.4 unless otherwise stated. To measure the concentration of free Ca^2+^ ions in the presence of AASTY or SMA2000 copolymers, copolymers were mixed with buffer for a final composition of 20 mM Hepes/NaOH, pH 7.4, 100 mM KCl and 0 mM, 1 mM, 3 mM or 7 mM CaCl_2_ and 0.1 % weight per volume (w/v) or 1 % copolymer. After 15 minutes incubation, ~10 μL buffer without copolymer was isolated using a Vivaspin-500 centrifugal concentrator with a 10 MW cutoff (Supp. Figure S3A). In a 96-well plate, 5 μL of this was mixed with 95 μL 0.13 mg/mL o-Cresolphthalein Complexone (oCPC) in 0.1 M CAPS, pH 10. The absorbance at 575 nm was measured on a Tecan Infinite M200 pro microplate reader and quantified based on a standard curve. To ensure that the copolymer had indeed been removed, spectra for select samples were measured using a NanoDrop1000 spectrophotometer.

### Determination of K_D_ between AASTY_7.4_-52 and Ca^2+^

Bound Ca^2+^ concentrations in the presence of 0, 0.05, 0.1, 0.25, 0.5, 0.75, 1, and 2 % AASTY_7.4_-52 and 7 mM CaCl_2_ were determined as described above by subtracting the measured free Ca^2+^ concentrations from the total Ca^2+^ amount present. The binding curve was fitted using SciPy’s^28^ curve-fit to the following equation, with the approximation that the all acrylic acid (AA) moieties can be considered free:

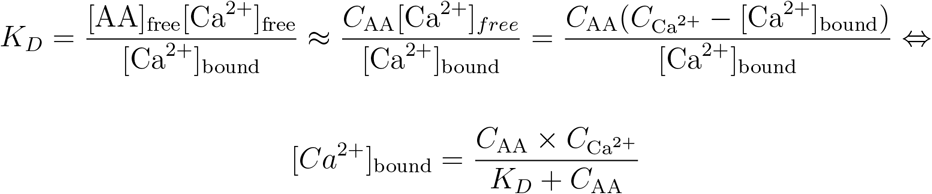

where the total AA concentration was estimated in the following manner from the mass fraction of AA in AASTY_7.4_-52, *w*_AA_, calculated based on the mole fractions of AA and styrene (STY), *x*_AA_ and *x*_STY_:

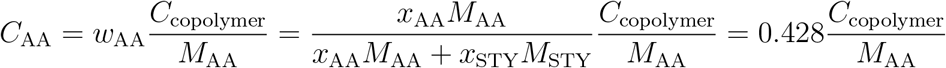

The resulting K*_D_* was verified by calculating free Ca^2+^ concentrations using the following equation, and comparing with those experimentally determined:

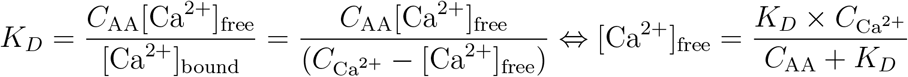

### Formation of empty nanodiscs

Similar to experiments previously described in Smith *et al*., ^19^ empty fluorescent nanodiscs were prepared by mixing 1 % copolymer with 1 mM lipids (small unilameller vecicles [SUVs] composed by 2 % fluorescent Lissamine Rhodamine B phosphatidylethanolamine [LissRhodPE, excitation at 554 nm and emission at 576 nm] and 98 % 1-palmitoyl-2-oleoyl-glycero-3-phosphocholine [POPC]) in 20 mM Hepes/NaOH and 100 mM KCl, unless otherwise specified in a 100 μL reaction volume. The mixture was incubated for 2 hours at 4°C before ultracentrifugation at 110,000 ×*g* and 4°C for 15 minutes. Free copolymer was removed by dialysis of the supernatant against buffer (three rounds of ~100 mL buffer per 100 μL sample) using dialysis tubing with a 100 kDa cut-off.

### Analysis by FSEC

Prior to analysis by FSEC, the sample was spin filtered through a 0.22 μm filter, to remove any remaining large aggregates. FSEC was run on a Superdex 200 increase column (5/150, Cytiva) attached to a Shimadzu (Shimadzu Europa GmbH) liquid chromatography system equipped with an autosampler (SIL-40), a fluorometer (RF-20A) and PDA detector (SPD-M40). The column was run at a flow rate of 0.2 mL/min in 20 mM Hepes/NaOH pH 7.4 and 100 mM KCl. 2 μL sample was loaded per run for pure copolymer samples (1 % or 0.1 %) and samples directly from solubilization. To assay effects of KCl and CaCl_2_ on formed nanodiscs, the nanodisc samples were diluted 10-fold to change the buffer. In this case, the 7.5 μL nanodisc sample was added to 67.5 μL buffer to reach the final specified concentrations. For dialyzed samples 20 μL of the diluted samples was loaded per run, while 10 μL were loaded for nanodisc samples that had not been dialyzed.

Most conditions were assayed with FSEC for a single run. However, a subset was tested multiple times either as different batches, i.e. experiments prepared on different days, or technical replicates as multiple injections from the same sample and on the same day to confirm reproducibility in the FSEC analysis (Supp. Figure S2). Because of the incomplete sampling of the repeats, error bars were left out of plots describing the stability of the free and nanodisc forming AASTY copolymer towards divalent ions in the main text figures.

### SEC data analysis

Size exclusion chromatography (SEC) data was analyzed using NumPy^29^ and SciPy.^28^ Areas under the curve were calculated using the composite trapezoidal rule through numpy.trapz() ^29^. To describe the absorption intensity, *I*(*V*), as a function of the elution volume, *V*, a sum of Gaussian functions was fitted using curve-fit from SciPy, using the following equation:

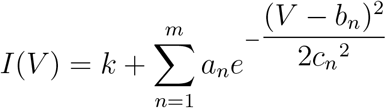

where *m* = 3 for AASTY_6.6_-44, AASTY_10_-47(-ttc), and SMA2000, while *m* = 4 for AASTY_8.9_-45 and AASTY_7.4_-52(-ttc), as three Gaussian curves did not yield a satisfactory fit for these samples. *k* is a constant offset to account for a possible background.

All plots were prepared with Matplotlib^30^ and schematics of workflows were created with BioRender.com.

## Results and Discussion

We previously introduced and characterized AASTY copolymers made with RAFT polymerization (Figure 1B), in terms of their ability solubilize different lipid compositions and a model mammalian membrane protein, in comparrison to SMA (Figure 1A).^19^ In that study, the AASTY copolymers had molecular weights in the range of 5-9 kDa as this range was reported effective for making nanodiscs with other copolymers. ^12^ In this work, we added an AASTY copolymer with a higher molecular weight and an intermediate acrylic acid content compared to those in Smith *et al*. ^19^ to test how these two parameters influence the Ca^2+^ sensitivity of the copolymer. RAFT polymerization of the AASTY copolymers result in a terminal Z group consisting of a trithiocarbonate (ttc) moiety that absorbs UV light with a max absorbance at 310 nm (Figure 1C). Assays were primarily conducted using polymers having the ttc group removed 1B), however for two of them, assays were also set up with the ttc-containing copolymers, as a secondary measure of copolymer stoichiometry in nanodisc formation. To better distinguish different AASTY copolymers as we expand the library in this and future studies, we here introduce a new naming convention, specifying the acrylic acid content (AA) and number-averaged molecular weight (M_n_): AASTY_Mn_-AA (Figure 1D). Hence, the new copolymer added in this work with a molecular weight of 12.5 kDa and acrylic acid content of 47 % is named AASTY_12.5_-47.

Characteristics of the AASTY copolymers used in this study are shown in Figure 1D.

### Acrylic acid content governs divalent cation sensitivity

Many proteins depend on Ca^2+^ and Mg^2+^ for their function. Some proteins are sensitive to the specific free Ca^2+^ concentration present, where too high concentrations are inactivating or desensitizing, ^31,32^ while complete absence of free Ca^2+^ can lead to irreversible inactivation. ^31,33^ Polyanionic copolymers such as AASTY are chelators of Ca^2+^ through their carboxylate anions. Consequently, the Ca^2+^ concentration that permits native nanodisc formation and stability is not necessarily equivalent to the amount of free Ca^2+^ ions required for a Ca^2+^ dependent protein to function, as the copolymer and protein are competing for Ca^2+^ ions. To shed light on this discrepancy, we used o-Cresolphthalein Complexone (oCPC) in a colorimetric detection of the available Ca^2+^ ions in a solution isolated after exposure to the AASTY copolymers (Figure 2A). Upon chelating divalent cations, oCPC yields a bright purple complex with an absorbance maximum at 575 nm. This allows a comparison of copolymers in terms of divalent cation binding, as the free ions left in solution will interact with oCPC (Figure 2B, Supp. Figure S3A).

**Figure 2:**
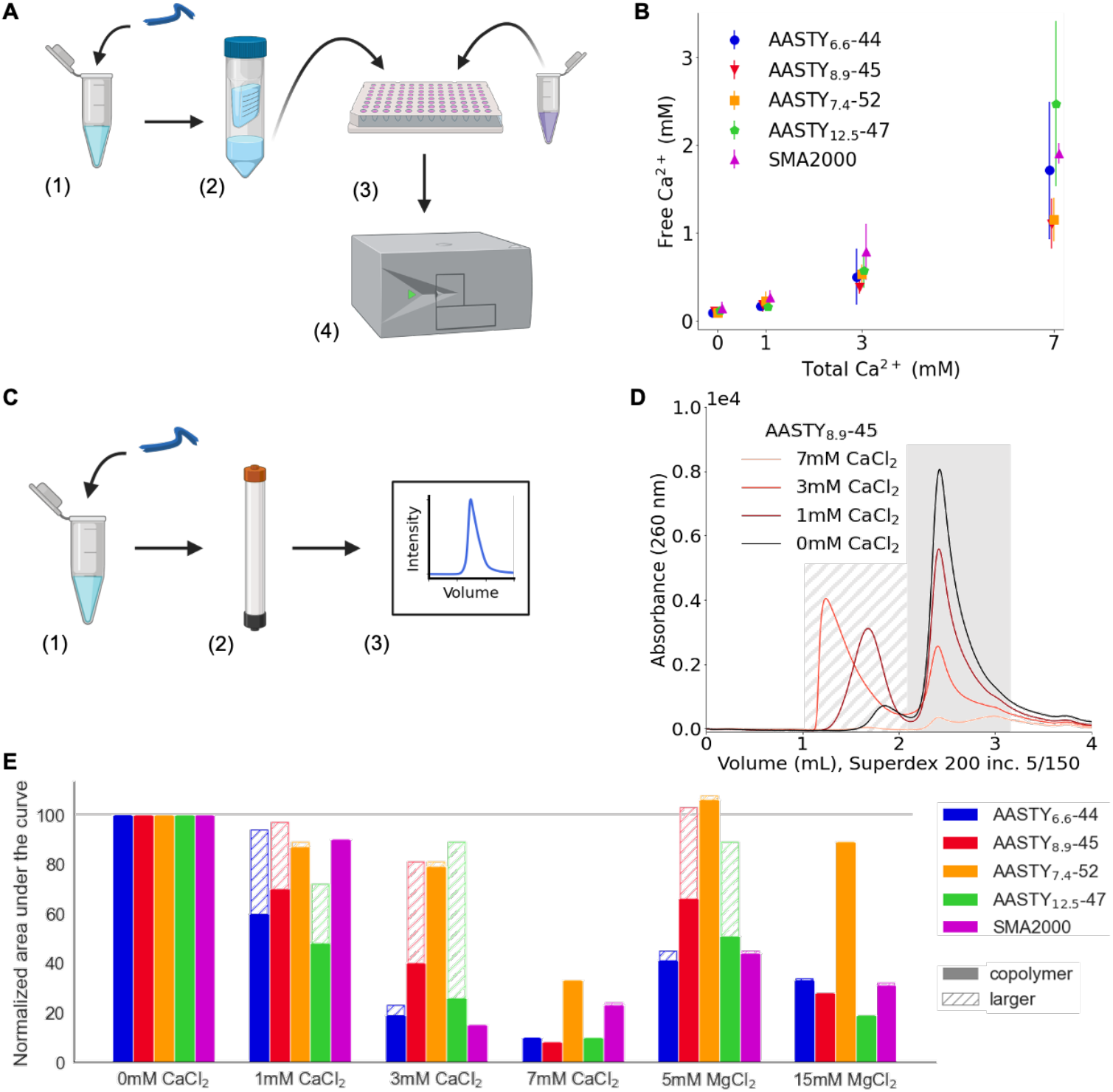
Interaction between free AASTY copolymer and the divalent cations Ca^2+^ and Mg^2+^. **A)** Free Ca^2+^ concentration was measured using oCPC in the following manner: Solubilized AASTY copolymer was mixed with buffer with or without CaCl_2_ (1). Next, buffer was isolated from the copolymer using a centrifugal concentrator with a 10 MW cutoff (2), and mixed with oCPC in a 96-well plate (3). Finally, the absorbance at 575 nm was measured using a plate reader (4). **B)** Measurements of free Ca^2+^ concentrations left in solution upon exposure to 1 % copolymer using oCPC. Each measurement was set up in triplicates. Small horizontal shifts have been introduced to separate the data points. Spectra of select samples can be seen in Supp. Figure S3A. **C)** Copolymers were analyzed by SEC by diluting them into a specified buffer (1) followed by loading on a Superdex 200 Increase 5/150 column running in 20 mM HEPES/NaOH, pH 7.4, and 100 mM KCl (2). The copolymer is observed at a wavelength of 260 nm (3, Supp. Figure S4). **D)** SEC of 0.1 % AASTY_8.9_-45 with different concentrations of CaCl_2_. The running buffer in all instances was without CaCl_2_. Chromatograms of other AASTY copolymers and with MgCl_2_ can be found in Supp. Figure S5. The shading indicates the species further quantified in E). **E)** Quantification of the traces from D) and Supp. Figure S5. Areas under the curve were determined separately for the copolymer peak (solid gray in D) and larger species (stripped grey in D) and normalized to the copolymer peak from 0 mM CaCl_2_ and MgCl_2_ for each copolymer individually. The grey line indicates 100 %. Each bar represents a single data point.

The assay with oCPC shows that the divalent cation binding capacity of the four AASTY copolymers included in this work only varies slightly (Figure 2B). Across all copolymers and assayed Ca^2+^ concentrations, only 21 % ± 6 % of the total Ca^2+^ ions remain free in the presence of 1 % copolymer as estimated from 2B. Thus, a higher absolute acrylic acid content is not equal to more binding of Ca^2+^ according to this assay.

The oCPC assay can also be used to determine the dissociation constant, K_D_, by titration of copolymer into a constant concentration of Ca^2+^. Thus, we determined the K_D_ of the carboxylate groups of AASTY_7.4_-52 to be 15 mM (Supp. Figure S3B). This K_D_ assumes that the copolymer is in excess leaving all potential Ca^2+^ sites free. This assumption is reasonable for high copolymer concentrations, but is not ideal for low copolymer concentrations at high Ca^2+^ concentrations where the copolymer is prone to precipitate. The K_D_ can be used to predict the free Ca^2+^ for a given concentration of the assayed copolymer and absolute Ca^2+^. It is worth noting that these predicted values correlate well with what is observed in the oCPC assay under similar conditions (Supp. Figure S3C).

The oCPC assay gauges an ensemble of particles in solution similar to other techniques typically used to characterize nanodisc forming copolymers. ^14,17,26^ In order to characterize the propensity of copolymers to aggregate in the presence of Ca^2+^ and Mg^2+^, we used SEC to separate and define the species present in solution. SEC was performed on copolymer solutions diluted into buffers containing different divalent cation concentrations (Figure 2C-E). The hydrophobicity of the copolymers vary, and we expected that this property would dominate their aggregation in response to divalent cations, an effect that would not be detected in the oCPC assay. For instance, AASTY_6.6_-44 has the largest content of styrene and is the most hydrophobic of the four AASTY copolymers (Figure 1D). Notably, due to the styrene moieties present in both AASTY and SMA, the copolymers absorb light in the UV range. Both have a maximum absorbance at 220 nm with decreasing absorbance at longer wavelengths (Supp. Figure S4B), with a local maximum at 260 nm for AASTY. However, differences in buffer composition between the sample (e.g. from ions) and running buffer during SEC will also show up at short wavelengths (Supp. Figure S4A), which is why we decided to use the absorbance at 260 nm (A_260_) to examine the behaviour of the AASTY copolymers in the presence of divalent cations (Figure 2D).

Upon addition of divalent cations, AASTY_6.6_-44, AASTY_8.9_-45, and AASTY_12.5_-47 formed species with smaller elution volumes (1-2 mL) indicative of formation or enrichment of larger, soluble species present before precipitation (Figure 2D and Supp. Figure S5A,B,D). In contrast, species at larger elution volumes (3 mL) were removed upon increased CaCl_2_. SMA2000 and AASTY_7.4_-52 precipitated from the solution at specific divalent cation concentrations, with limited formation of soluble aggregates (Supp. Figure S5C,E). The observations from the SEC runs of the four AASTY copolymers and SMA2000 are summarized and quantified in Figure 2E. This indicates that at least three of the AASTY copolymers have a “sweet spot”, i.e. a divalent ion concentration at which it forms larger species, before it precipitates from the solution: While 3 mM and higher CaCl_2_ concentrations removed the majority of AASTY_6.6_-44 and SMA2000 from solution, AASTY_8.9_-45 and AASTY_12.5_-47 remaind in solution at this concentration. AASTY_7.4_-52 was only significantly removed from solution, presumably due to aggregation, in the presence of 7 mM CaCl_2_, at which all other AASTY copolymers are also largely precipitated. The SEC experiment highlights the importance of a thorough characterization of the divalent cation tolerance of nanodisc forming copolymers, as it can be overestimated due to soluble aggregates. A similar trend is observed upon addition of Mg^2+^, though the tolerance is higher for all five assayed copolymers. Here, soluble aggregates are observed for AASTY_8.9_-45 and AASTY_12.5_-47 at 5 mM MgCl_2_, while only AASTY_7.4_-52 remains significantly in solution at 15 mM MgCl_2_ (Figure 2E).

In context of the observed Ca^2+^ binding capacities from the oCPC assay (Figure 2B), we see that while the copolymers have comparable binding of Ca^2+^, the AASTY copolymers with a low acrylic acid content AASTY_6.6_-44, AASTY_12.5_-47 and SMA2000 are the most sensitive to precipitation in the presence of Ca^2+^. Moreover, the copolymer with the highest acrylic acid content, AASTY_7.4_-52, is the least sensitive to divalent cations. This suggest that more carboxylate negative charge shield these copolymers from aggregation, allowing AASTY_7.4_-52 to withstand a higher level of Ca^2+^ ions before precipitating.Additionally, a higher molecular weight also decreases divalent cation tolerance, as it is seen that AASTY_8.9_-45 remains in solution at concentrations of Ca^2+^ where AASTY_12.5_-47 forms large species and precipitates. Altogether, precipitation is governed by molecular weight and charge, with increasing charge conveying stability, and increasing molecular promoting precipitation. We next sought to test whether this effect also holds true when polymers participate in nanodisc formation.

### Divalent cations influence nanodisc formation

As apparent from the previous section, the copolymers are able to generate larger species in the presence of divalent cations. In some experimental setups, these larger species could be confused for nanodiscs (Figure 2D). Therefore, to be able to monitor the behaviour of nanodiscs and separate them from any free copolymer present in the solution, we generated fluorescent nanodiscs by solubilizing 1 mM SUVs composed by 98 % POPC and 2 % of fluorescent LissRhodPE with 1 % copolymer (Figure 3A-C). In our previous study of AASTY,^19^ we found that the efficiency of the nanodisc formation with the AASTY copolymers depends on the lipid composition and charge. As three of the four AASTY copolymers solubilized the zwitterionic synthetic lipid POPC into nanodiscs exceedingly well, we decided to work exclusively with POPC as the solubilizing lipid and the fluorescent lipid LisRhodPE as our reporting lipid in this study as well. We note that divalent cations are expected to interact with both type of lipids to some extent.

**Figure 3:**
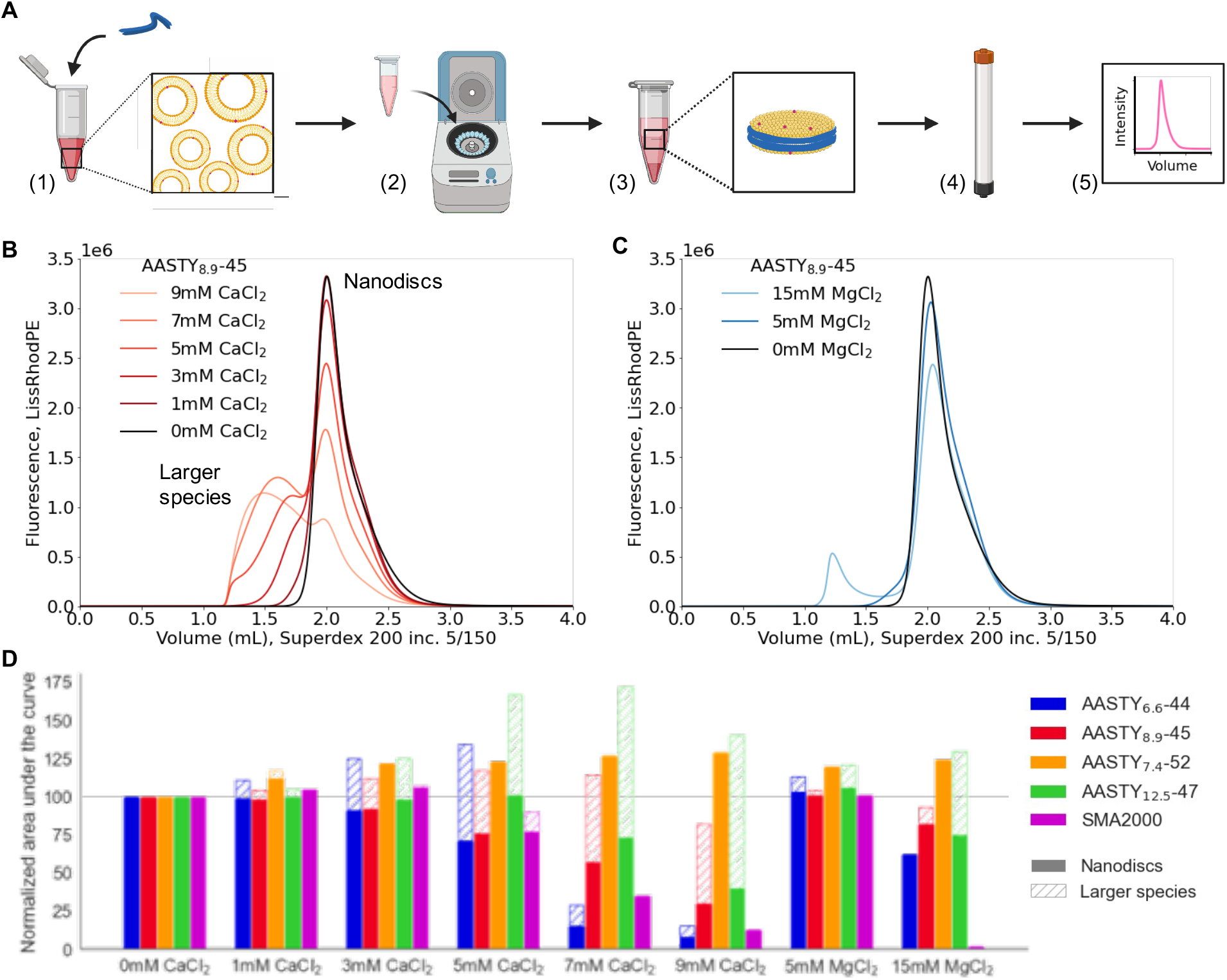
Solubilization of SUVs in the presence of divalent cations. **A)** Fluorescent nanodiscs are produced in the following manner: SUVs of 98 % POPC and 2 % fluorescent LissRhodPE are incubated with copolymer (1) followed by ultracentrifugation (2) to remove insoluble material. The supernatant containing fluorescent nanodiscs and free copolymer (3) is analyzed by FSEC (4,5). **B)** FSEC profiles of nanodiscs generated by solubilizing 1 mM lipids with 1 % AASTY_8.9_-45 in the presence of specified CaCl_2_ concentrations. **C)** Same as B) with MgCl_2_. Profiles for the other copolymers can be found in Supp. Figure S7. **D)** Quantification of the area under the curve for the nanodisc peak (at 2 mL) and larger species normalized to the nanodisc peak at 0 mM divalent cations. The nanodisc peak in each trace was determined as the part that falls within the 0 mM trace of that copolymer. The grey line indicates 100 %. A total fluorescence higher than this, could be due to hydrophobic effects increasing the fluorescence from partially aggregated species. Each bar represents a single data point.

Subjecting the nanodiscs to FSEC in the absence of divalent cations, we observe that the copolymers generate nanodiscs with peaks of varying elution volumes, which we interpret as the nanodiscs being different in size (Supp. Figure S6). Curiously, the elution volume or the size of the nanodiscs does not correlate with the molecular weight of the copolymer (Supp. Figure S6C,E): While AASTY_6.6_-44, AASTY_8.9_-45, and AASTY_12.5_-47 produce discs of very similar sizes, the discs produced by AASTY_7.4_-52 and SMA2000 are larger. In fact, the FSEC profiles for the AASTY copolymers support that the nanodisc size is affected by the composition, i.e. the carboxylic acid content, while the molecular weight is inconsequential (Supp. Figure S6D,E).

Some proteins require a constant presence of calcium and magnesium to retain activity. ^31^ Therefore, we next sought to use FSEC to characterize which CaCl_2_ and MgCl_2_ concentrations are permissible during nanodisc formation with the four AASTY copolymers and SMA2000 (Figure 3B-D and Supp. Figure S7). As with the copolymer in the absence of lipids, Ca^2+^ ions have a more pronounced effect on the elution volumes and presumably the sizes of the nanodiscs than Mg^2+^ ions for all assayed copolymers (Figure 2E and Figure 3D). Similar to the free copolymer, the proportion of larger, soluble species increases in the nanodisc samples as the CaCl_2_ concentrations increase. This comes at a cost of well-formed nanodiscs. An explanation could be the rouleaux stacking of discs observed by Ravula *et al*. under similar conditions.^27^ Notably, AASTY_7.4_-52 is least affected by the presence of divalent cations both in nanodiscs and in its free form, and while all nanodiscs tolerate the presence of 3 mM CaCl_2_ reasonably well, their behaviours start to diverge significantly beyond this concentration (Figure 3D). For instance, the AASTY_12.5_-47 nanodisc peak broadens at 5 mM CaCl_2_ and shifts to larger size at 7 mM and 9 mM CaCl_2_, but remains in solution (Supp. Figure S7E). AASTY_6.6_-44 and AASTY_8.9_-45 nanodiscs both form large, soluble species before precipitating, though AASTY_8.9_-45 does so at higher CaCl_2_ concentrations. In contrast to the AASTY polymers, SMA2000 precipitates to large extent. In terms of stability towards Mg^2+^, all five copolymers tolerates 5 mM MgCl_2_, while SMA2000 nanodiscs are unstable in 15 mM MgCl_2_. In conclusion, comparing the four types of AASTY nanodiscs, the tolerance towards CaCl_2_ and MgCl_2_ is dependent on the copolymer composition and molecular weight. Our FSEC data support that the large carboxylic acid content in AASTY_7.4_-52 helps stabilize it in nanodiscs, while the tendency of AASTY_12.5_-47 to form soluble aggregates is caused by its larger molecular weight (Figure 3D).

### AASTY is less tolerant to Ca^2+^ when in nanodiscs than as unbound copolymer

Next, we wanted to decipher whether the copolymers are less sensitive towards Ca^2+^ ions when incorporated into a nanodisc than when pure in solution. At a first glance, this seems to be the case (compare Figure 3D with Figure 2E). However, this observation is an effect of the higher copolymer concentrations present during lipid solubilization (1 %), compared with the SEC experiments with the pure copolymers (0.1 %, see Experimental). Looking at the A_260_ SEC traces of the AASTY nanodisc samples revealed that there is a large amount of free, excess copolymer in the solution (Figure 4A, insert). Thus, we used the absorbance at 260 nm to assess the behaviour of the excess AASTY not part of nanodiscs when in presence of Ca^2+^. Comparing nanodiscs diluted to a total copolymer concentration of 0.1 % in buffer with CaCl_2_, we see that AASTY incorporated into nanodiscs is less resistant to Ca^2+^ than the free, excess AASTY copolymer (Figure 4). While the free AASTY copolymer in a nanodisc sample have similar elution profiles as the pure copolymer in the absence of lipids at varying CaCl_2_ concentrations, the nanodiscs aggregate and precipitate at lower CaCl_2_ concentrations. This is most pronounced for AASTY_8.9_-45 where 1 mM CaCl_2_ is enough to destabilize the discs, while the difference only becomes apparent for AASTY_7.4_-52 at 7 mM CaCl_2_, where no nanodiscs are left in solution, while ~35 % of the free copolymer remains in solution. It should be noted that the apparent stability of AASTY_8.9_-45 nanodiscs at 3 mM CaCl_2_ is an artefact of the quantification, as the nanodisc peak has in fact shifted entirely to smaller elution volumes and hence larger size at this point (see Supp. Figure S8A). Lipid-associated copolymer in nanodiscs has reduced degrees of freedom compared to the free copolymer, which could cause the nanodiscs to stick together and aggregate at lower Ca^2+^ concentrations, while divalent cation binding to the lipid headgroups could cause rouleaux stacking of nanodiscs as observed by Ravula *et al*.. by NMR^27^

**Figure 4:**
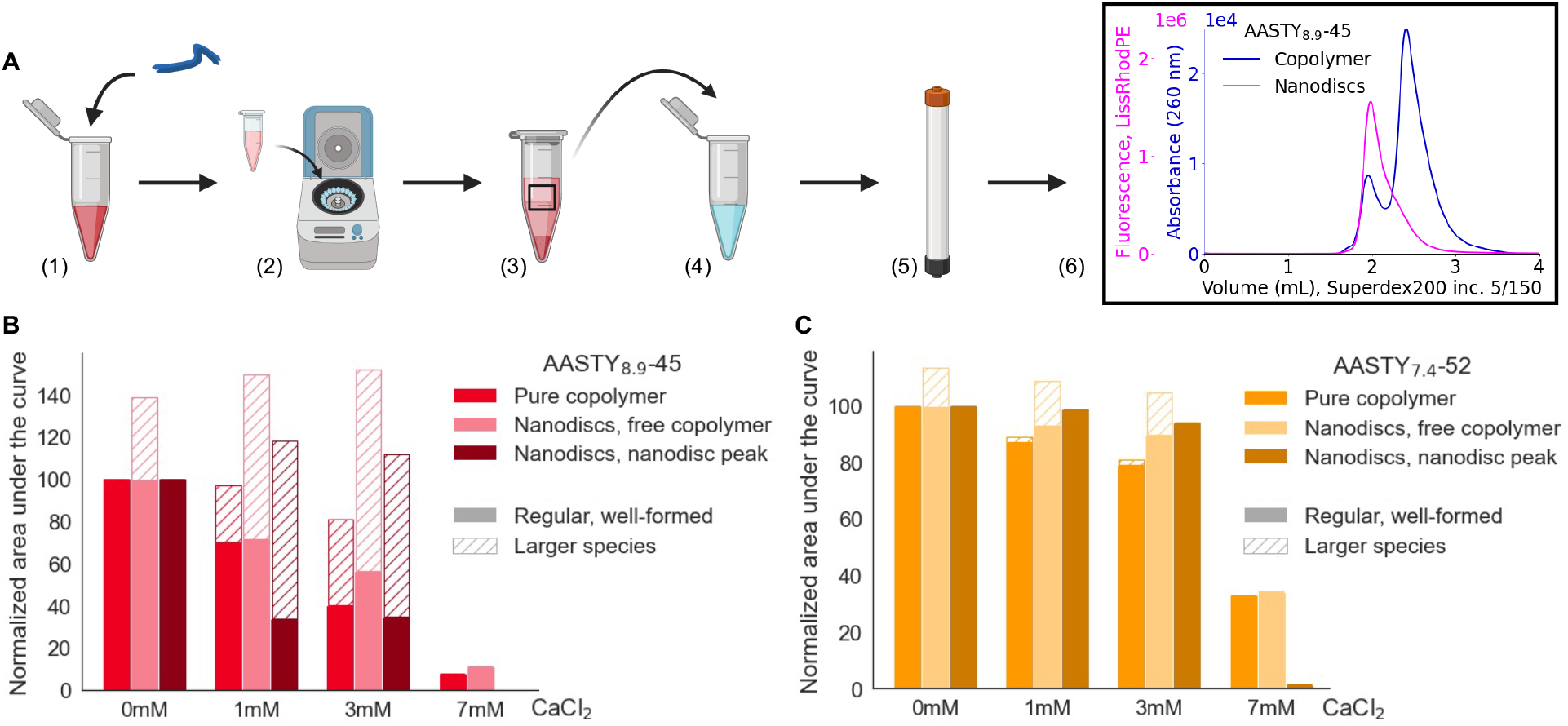
Comparison of Ca^2+^ resistance of AASTY copolymers in absence of lipids (pure form) and in presence of lipids (nanodiscs and free, excess form). **A)** After generation of fluorescent nanodiscs (1-3, as in Figure 3A), the supernatant containing the nanodiscs was diluted 10-fold into buffer with specified concentration of CaCl_2_ (4), before analysis by UV and fluorescent SEC (5,6). The insert shows how A_260_ and fluorescent SEC traces were used to monitor and compare the elution profiles of the styrene-containing AASTY copolymers (blue) and the fluorescent AASTY nanodiscs (pink). **B)** Quantification of area under the curve for loading of pure AASTY_8.9_-45 (as in Figure 2E), compared with the area under the curve of the free copolymer peak at A_260_ for diluted AASTY_8.9_-45 nanodiscs (elution volume of ~2.5 mL) and the area under the curve for the nanodisc peak seen by fluorescence of LissRhodPE (as in Figure 3B). For the free copolymer from the nanodisc sample, the larger species also include nanodiscs. A_260_ and fluorescent SEC traces of the diluted AASTY_8.9_-45 nanodisc samples are included in Supp. Figure S8A,B. In all instances, the running buffer was 20 mM Hepes/NaOH pH 7.4, 100 mM KCl without CaCl_2_. Each bar represents a single data point. **C)** Same as B), but for nanodiscs prepared with AASTY_7.4_-52. A_260_ and fluorescent SEC traces of the diluted AASTY_7.4_-52 nanodiscs are included in Supp. Figure S9A,B.

The higher divalent cation sensitivity of lower copolymer concentrations, aligns well with our observation that a lower copolymer concentration results in higher free Ca^2+^ concentrations (Supp. Figure S3B), meaning that some Ca^2+^ binding by the copolymer can protect it against precipitation. Taken together, our SEC data of AASTY nanodisc formation in the presence of CaCl_2_ and MgCl_2_ show that copolymer resistance towards divalent cations is highly dependent on the assayed copolymer concentration and highlight that a single, limiting Ca^2+^ concentration for a given copolymer is insufficient as a guideline. Our data also shows that the common practice of using turbidity as a measure of divalent cation tolerance can be misleading for describing copolymer lipid nanodisc tolerance of divalent cations. Our data show that free, excess copolymer can function as a cushion to the effects of divalent cations on the system, by binding ions and forming various soluble species without changes in turbidity. Furthermore, the lower divalent cation tolerance of lipid-associated copolymer can be masked by the presence of free copolymer with higher tolerance.

### Dialysis removes free copolymer and affects size of discs

As we hypothesized that the excess, free copolymer present after nanodisc formation is able to act as a cation sink, we decided to examine how the AASTY nanodiscs behave in the presence of CaCl_2_ upon removal of the excess copolymer.^34,35^ Thus, we subjected the nanodisc samples to dialysis (Figure 5). During a protein purification, this excess copolymer would typically be removed by an affinity step. Here, we used a membrane with a 100 kDa cut-off, which is sufficient to retain the nanodiscs. Progression of the dialysis was monitered with FSEC, with the free copolymer peak decreasing in size as dialysis progressed. It should be noted that dialysis is not feasible for SMA due to its high dispersity.

**Figure 5:**
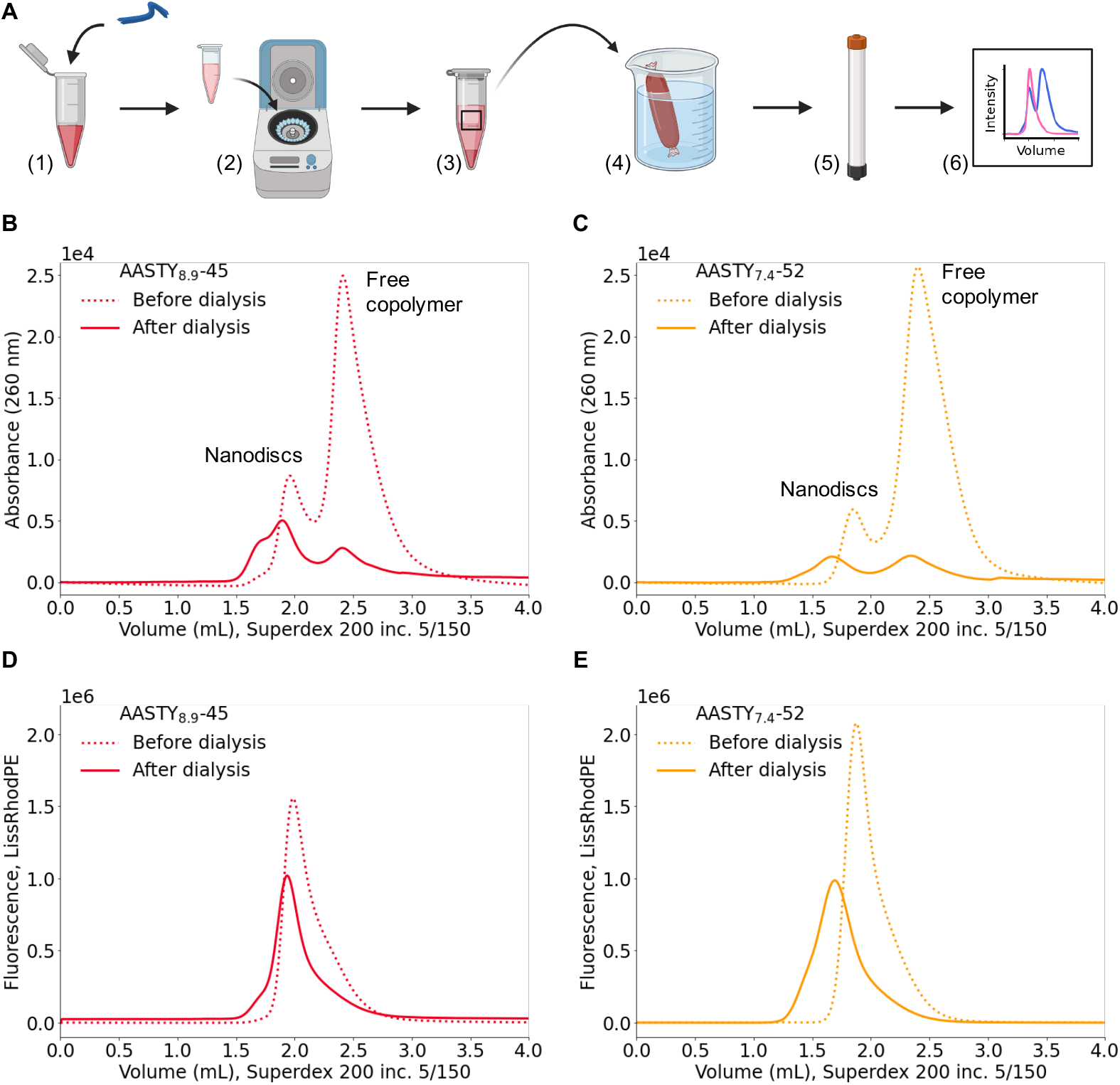
Removal of free copolymer by dialysis. **A)** After generation of fluorescent nanodiscs (1-3, as in Figure 3A), the supernatant is dialyzed against a large excess of buffer to remove free copolymer (4), before analysis by FSEC (5,6). Visualization of **B)** AASTY_8.9_-45 and **C)** AASTY_7.4_-52 nanodiscs and free copolymer by absorbance at 260 nm before and and after dialysis. FSEC visualizing the **D)** AASTY_8.9_-45 and **E)** AASTY_7.4_-52 nanodiscs with fluorescent lipids before and after dialysis. The dialysis resulted in 2-3-fold dilution of the sample.

In addition to removing free, excess copolymer, dialysis of the nanodiscs imposes a shift of the nanodisc peak to slightly lower elution volumes for both AASTY_8.9_-45 and AASTY_7.4_-52, with broadening of the AASTY_7.4_-52 nanodisc peak (Figure 5D,E), suggesting that the discs become larger and more disperse in size. This fits with the altered lipid:polymer ratio of these samples, which has also been observed for SMA.^36^ Our findings corroborate the study by Mauro *et al*. who showed that removing excess polymer resulted in an increased critical micelle concentration for DOPC copolymer mixtures. ^37^

A comparison of the nanodisc sample prior to dialysis (Supp. Figure S8A and S9A) with the dialyzed sample (Supp. Figure S8C and S9C), shows that the dialysis and hence the removal of free, excess copolymer influences the FSEC traces. Between the two assayed nanodisc forming copolymers, AASTY_8.9_-45 and AASTY_7.4_-52, the differences are particularly striking for the latter: For AASTY_7.4_-52, the nanodisc sample prior to dialysis is stable at CaCl_2_ concentrations up to 5 mM (elution volumes around 2 mL), while a large proportion of the dialyzed nanodiscs shift to the void elution volumes around 1.25 mL already in the presence of 1 mM CaCl_2_ (compare Supp. Figure S9A with S9C). This indicates that it is likely calcium binding by the free AASTY_7.4_-52 copolymer that is responsible for the stability of the nanodisc sample towards CaCl_2_ concentrations up to 5 mM. When the free copolymer concentration is reduced upon dialysis, higher free Ca^2+^ concentrations are present than in the undialyzed sample according to Supp. Figure S3B.

### KCl stabilizes AASTY nanodiscs in presence of CaCl_2_ and absence of free copolymer

It has been shown that salts and positively charged amino acids improve affinity chromatography purification of membrane proteins in native nanodiscs. ^19,22,23^ Furthermore, benefits of high ionic strengths on nanodisc formation has been described for SMA^38,39^ and DIBMA.^40^ We hypothesized, that monovalent salts such as KCl would also lead to an improved tolerance of AASTY nanodiscs towards Ca^2+^. To assess this, the dialyzed AASTY_8.9_-45 and AASTY_7.4_-52 nanodisc samples were diluted 10-fold into buffer with various CaCl_2_ and KCl concentrations and subjected to A_260_ and fluorescent SEC analysis (Figure 6A). The FSEC traces show, that KCl shifts the nanodiscs peak towards larger elution volumes, indicating a smaller size closer to the nanodisc size prior to dialysis (Figure 5D,E and Figure 6B,D). It is worth noting that the nanodisc peak is quite broad for AASTY_7.4_-52 in 100 mM KCl and 0 mM CaCl_2_ (Figure 6D, solid black). Addition of 2 mM CaCl_2_, while producing a large void peak, also gives a nanodisc peak, which elutes at slightly lower size and appears sharper (Figure 6D, solid red), suggesting that a higher ionic strength in the buffer is necessary for AASTY_7.4_-52 nanodiscs to be stable even in the absence of divalent cations. The FSEC data is quantified in Figure 6C,E and shows that adding 300 mM and 700 mM KCl ameliorates aggregation and precipitation of the assayed AASTY nanodiscs in the presence of increasing Ca^2+^. For AASTY_8.9_-45, 700 mM KCl is notably better for Ca^2+^ tolerance than 300 mM (Figure 6C), while 700 mM KCl only improves the nanodisc peak for AASTY_7.4_-52 marginally over 300 mM KCl (Figure 6E). Taken together, our data corroborates with the observed benefits of adding salts or amino acids to copolymer nanodisc preparations, and extends this to protection against Ca^2+^.^19,22,23,38-40^ Evidently, it is beneficial to screen salt concentrations to fine-tune the sample composition and stability for a specific purpose for a particular membrane protein which, like the copolymers, also has a net charge.

**Figure 6:**
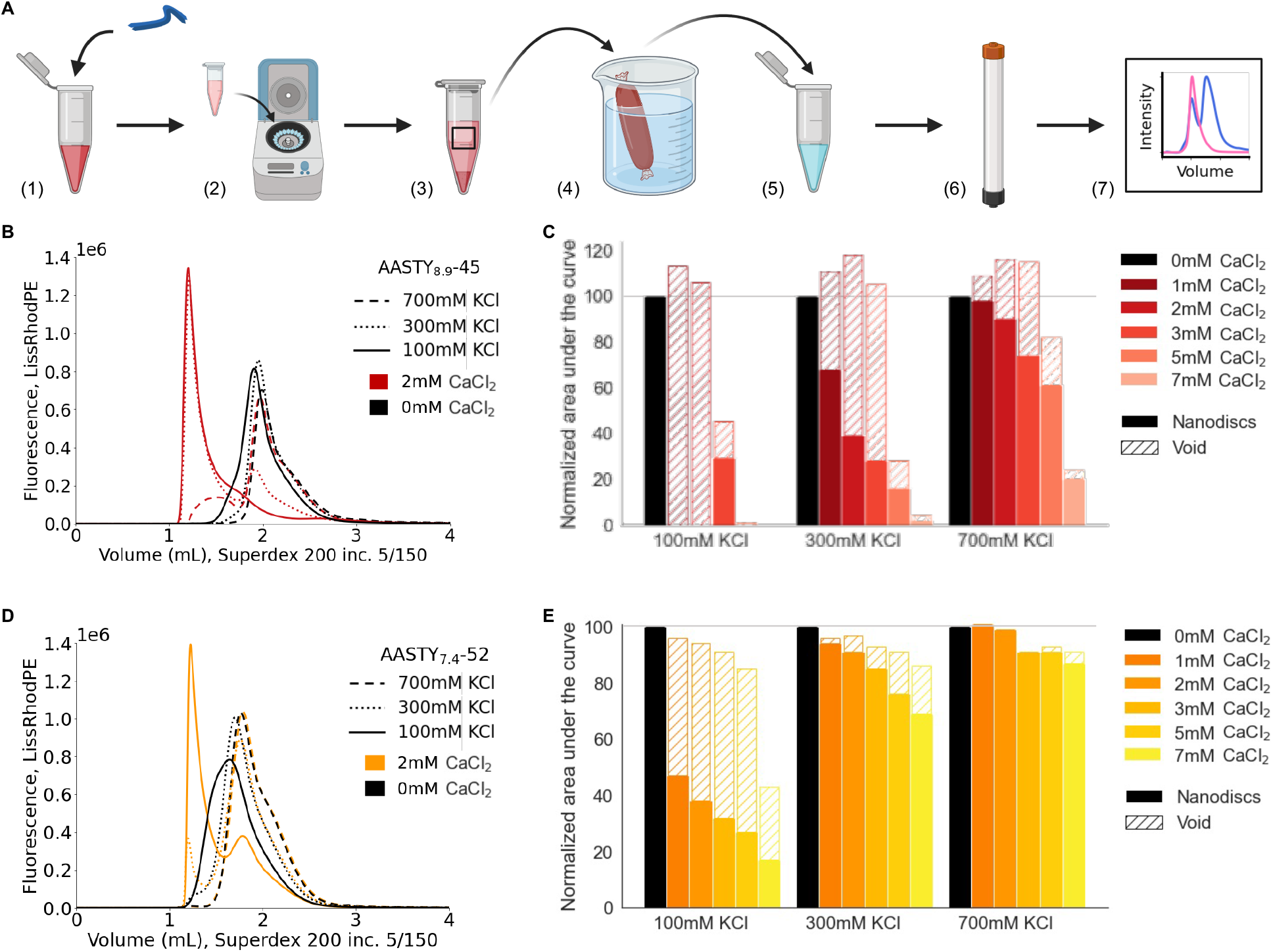
Stability of dialyzed nanodiscs in the presence of Ca^2+^ ions at different ionic strengths. **A)** After removal of free copolymer from a fluorescent nanodisc sample (steps 1-4, as in Figure 5A), the discs were diluted into buffers with varying KCl and CaCl_2_ concentration (5), before analysis by FSEC (6,7). **B)** Select FSEC traces of fluorescent AASTY_8.9_-45 nanodiscs at different KCl concentrations with or without 2 mM CaCl_2_, prepared by 10-fold dilution of the dialyzed sample from Figure 5B,D into specified buffer. The running buffer in all instances was 20 mM Hepes/NaOH pH 7.4, 100 mM KCl without CaCl_2_. Additional CaCl_2_ concentrations can be found in Supp. Figure S8. **C)** Quantification of the area under the curve for traces from B) and Supp. Figure S8C,E,G divided into void and nanodisc peaks. For each KCl concentration separately, the values were normalized to the 0 mM CaCl_2_ trace. Each bar represents a single data point. **D,E)** Same as B,C) but for nanodiscs prepared with AASTY_7.4_-52 (Figure 5C,E). Complete traces for dilutions of the AASTY_7.4_-52 nanodiscs are found in Supp. Figure S9.

### Nanodisc size is dynamic and depends on lipid concentration

Evidently, the thermodynamics of nanodisc formation are very complex: Extrinsic factors such as the ionic strength of the environment and the copolymer concentration affect the apparent size of the nanodiscs (Figure 5B,D and 6). We previously observed that the lipid composition and charge influence AASTY nanodisc formation. ^19^ Furthermore, the process of nanodisc formation is likely also affected by the membrane proteins that one wish to incorporate into the nanodiscs. However, intrinsic properties of the individual copolymers also play a role (Supp. Figure S6).

To look further into determinants of nanodisc size, we examined the effects of different lipid concentrations on the size of resulting nanodiscs as quantified by the elution volumes and area under the curve of the resulting fluorescent and A_260_ SEC traces (Figure 7). According to our data, higher lipid concentrations lead to lower elution volumes and hence larger nanodiscs (Figure 7D). This observation corroborates with the larger nanodiscs observed after dialysis (Figure 5B,D), as the lipid:polymer ratio is increased in experiments with both AASTY_8.9_-45 and AASTY_7.4_-52. Our analysis shows that AASTY nanodiscs are not static once formed; indeed mixing the largest and smallest nanodiscs leads to nanodiscs of intermediate size, coinciding perfectly with their resulting lipid concentration (Figure 7A,C). Our observations of the dynamic nanodiscs corroborates with previous findings of lipid exchange in SMA and DIBMA nanodiscs,^35,41^ for which a dependence of nanodisc size on the lipid:polymer ratio was also reported. ^11,36^

**Figure 7:**
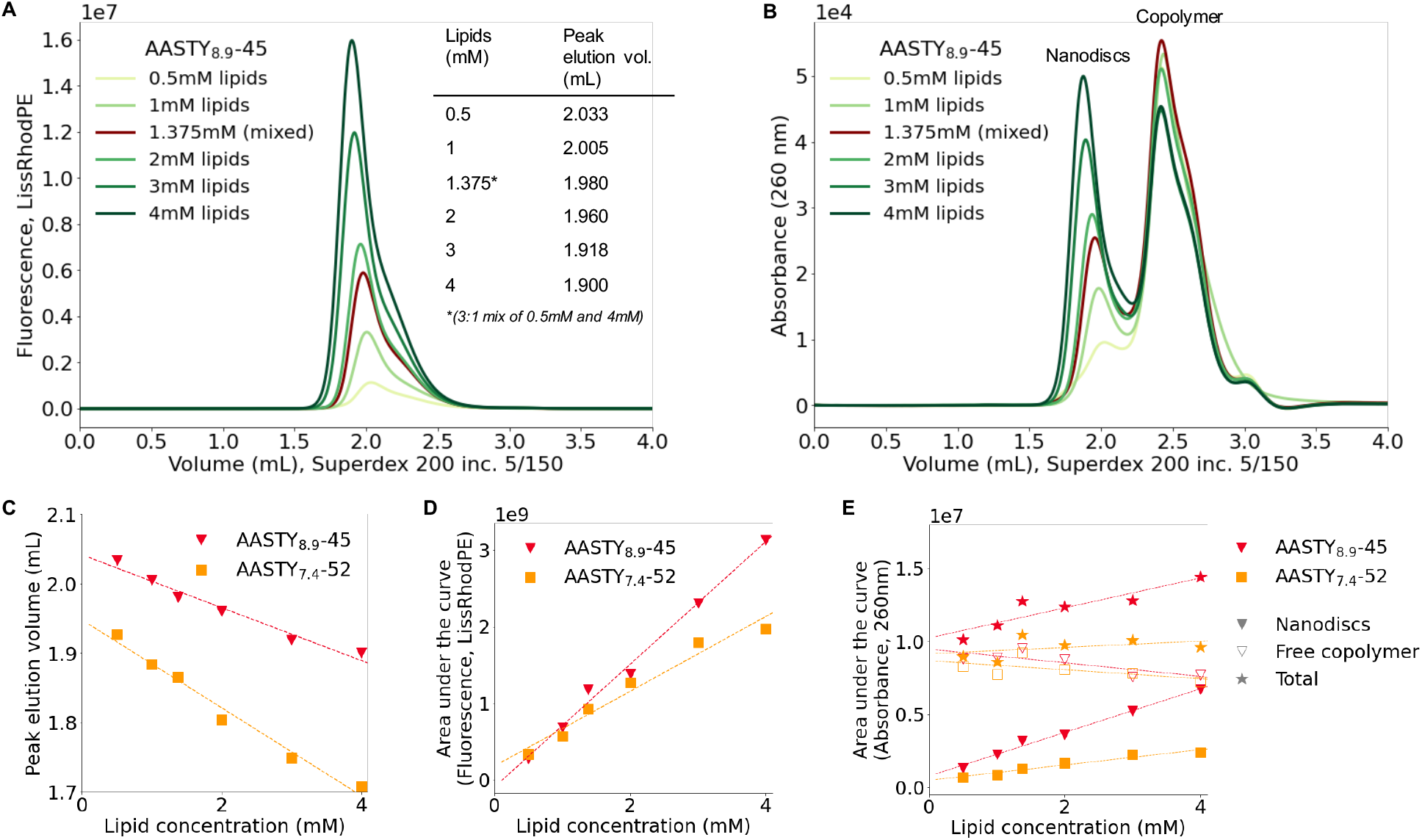
The effect of lipid concentration on nanodiscs size. **A)** FSEC traces of nanodiscs generated by solubilizing different total lipid concentrations (98 % POPC and 2 % Liss-RhodPE) with 1 % AASTY_8.9_-45. The table insert indicates the elution volume of the peak for different samples. The sample with 1.375 mM lipids (brown trace) was produced by mixing the 0.5 mM and 4 mM samples. **B)** A_260_ SEC trances for the samples in A). While the nanodisc peak shifts to lower elution volume, the free copolymer peak remains in the same position. **C)** The elution volume of the peak fractions plotted as a function of the lipid concentration. Lower elution volume translates to larger nanodiscs. **D)** Area under the curve from the fluorescent nanodisc signal as a function of the lipid concentration. **E)** Area under the curve from A_260_ for the nanodisc and copolymer peaks separately as well as the total. Plots analogous to A,B for AASTY_7.4_-52 can be found in Supp. Figure S10.

Increasing the lipid concentrations, increases the area under the curve in both the fluorescent and A_260_ SEC traces (Figure 7D,E). Interestingly, the size of copolymer peak originating from the the free copolymer not incorporated into nanodiscs changes little with the amount of lipids (Figure 7B,E). Thus, it seems like the total absorbance of the nanodisc sample increases although the copolymer concentration remains constant. This discrepancy shows that it is not straightforward to determine the fraction of copolymer that goes into nanodiscs as the spectral properties of the free, unbound copolymer, appear different from the copolymer forming nanodiscs. This difference likely stems from the different chemical environments experienced by styrene. The difference is clear when comparing scaled chromatograms from different wavelengths (Supp. Figure S11A and S12A for AASTY_7.4_-52 and AASTY_12.5_-47).

As an attempt to account for individual AASTY copolymers, we utilized copolymers with the ttc end group intact from the RAFT synthesis (Figure 1C) in solubilization experiments for AASTY_7.4_-52 and AASTY_12.5_-47. ttc absorbs light at 310 nm, and thereby functions as a secondary chromophore to styrene for estimating the distribution of copolymer across different species. As each copolymer contains exactly one ttc, the signal should be proportional to the number of copolymer molecules.

To quantify the fraction of free copolymers, Gaussian functions were fitted to the chromatogram (Figure 8A,C). This approach was particularly necessary for AASTY_12.5_-47, where the nanodisc peak overlaps with that of the free copolymer (Figure 8C). Calculating the fraction of free copolymer, we observe that the value obtained across different wavelengths is fairly stable for AASTY_7.4_-52 and AASTY_7.4_-52-ttc with 84 % and 82 % of the copolymer population estimated to be free with 1 mM lipids solubilized (Figure 8B). For AASTY_12.5_-47-ttc on the other hand, 87 % is estimated to be free based on wavelengths where styrene absorbs strongly, while 78 % is estimated from those where ttc absorption is dominating (Figure 8D). The overestimation of free copolymers from the A_260_ suggests that styrene absorbs less when engaged in interactions with lipids. However, it cannot be ruled out that the absorbance of ttc is also affected by association with lipids. Using the free copolymer fraction to quantify nanodisc participation, we find a decrease in free copolymer with a higher solubilized lipid concentration (Figure 8B,D and Supp. Figure S13). As such, we see that a larger fraction of copolymer is engaged in nanodiscs when more lipids are solubilized, though the majority remains free in all tested conditions (Supp. Figure S13). In total, both methods of quantification generally corroborate the picture that the majority of copolymers remain free when making nanodiscs.

**Figure 8:**
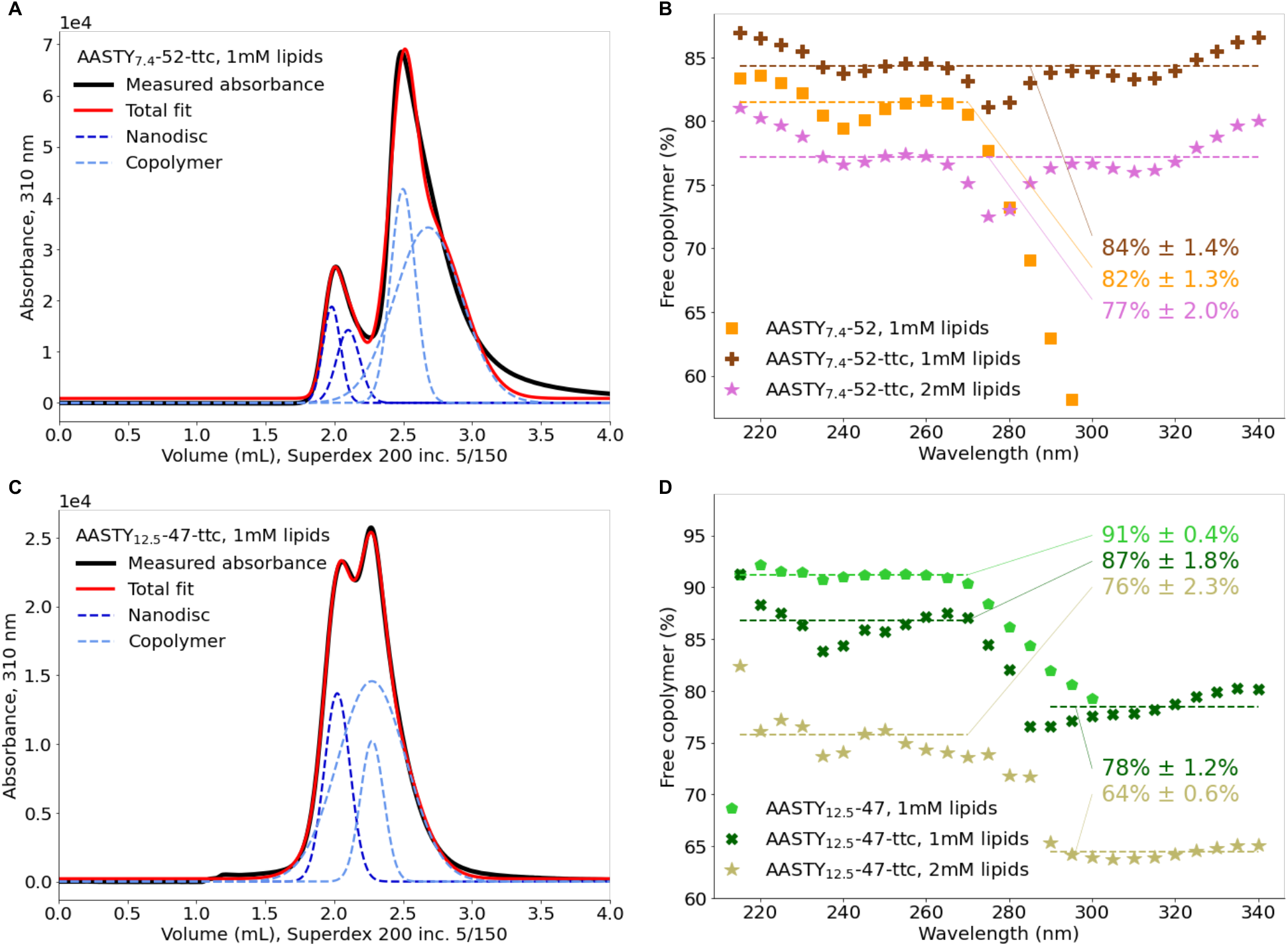
Quantification of free copolymer after solubilization of lipids. **A)** Absorbance at 310 nm for 1 mM lipids (98 % POPC and 2 % LissRhodPE) solubilized with AASTY_7.4_-52-ttc with four Gaussian functions fitted to describe the measured absorbance. Two are attributed to nanodiscs and two to the copolymer. The normalized spectrum of AASTY_7.4_-52-ttc can be found in Supp. Figure S11B,C. **B)** The fraction of free copolymer estimated from the area under the curve of the Gaussian functions describing the copolymer (as shown in panel A) at different wavelengths for AASTY_7.4_-52 and AASTY_7.4_-52-ttc after solubilization of specified lipid concentrations. The average value is indicated on the plot and shown as a dashed line. There is very little signal for AASTY_7.4_-52 at wavelengths longer than 280 nm, this average is only calculated from wavelengths 215-270 nm. Absorbance profiles at 260 nm for all three samples and the total area under the curve are shown in Supp. Figure S11D,E. **C,D)** Analogous to A,B), but for AASTY_12.5_-47 and AASTY_12.5_-47-ttc. Here, only three Gaussian functions were used to describe the data, where one was attributed to nanodiscs. In D) two different averages were calculated for AASTY_12.5_-47-ttc, as wavelengths dominated by the copolymer and the ttc tag gave very different estimates. Additional data for AASTY_12.5_-47-ttc is shown in Supp. Figure S12.

## Conclusions

In this work, we characterize the effects of Ca^2+^ and Mg^2+^ on the stability of AASTY copolymers and nanodiscs using a coupled dye assay and analytical UV and fluorescence SEC. We find that the copolymer composition and molecular weight, have a significant effect on its propensity to aggregate and precipitate when exposed to divalent cations. We observe that a higher acrylic acid content conveys apparent tolerance of AASTY for divalent cations, though this tolerance is also dependent on copolymer concentration, and excess, free copolymer.

Our data shows that the AASTY copolymers tolerate higher concentrations of Mg^2+^ than Ca^2+^, both in nanodiscs and in absence of lipids. In the presence of 100 mM KCl, AASTY copolymers form large, soluble aggregates and precipitates in the presence of increasing concentrations of divalent cations. This instability is exacerbated if the free, excess AASTY is removed by dialysis and ameliorated by increasing the ionic strength of the buffer. We show that AASTY copolymer lipid nanodiscs tolerate concentrations up to at least 7 mM CaCl_2_ when in the presence of 700 mM KCl.

Quantification of the free copolymer from the SEC data shows that increasing the lipid concentrations engage more copolymer into disc formation, while also forming larger nanodiscs. However, the majority of the added copolymers remain free in all assayed conditions, varying from 65% to 90%, depending on the method of quantification and the polymer in question.

At a concentration of 1 %, all four AASTY copolymers as well as SMA2000 bind approximately 80 % of the present Ca^2+^ ions. As such, only a fraction of the present Mg^2+^ and Ca^2+^ ions would be accessible to a potential membrane protein of interest, and this concentration would change throughout protein purification when the copolymer concentration changes. These effects are essential to be aware of when working with Mg^2+^ or Ca^2+^ sensitive membrane proteins in native nanodiscs.

## Supporting Information Available

Supporting information contains the following supplementary figures: NMR spectrum of AASTY_8.9_-45 and AASTY_12.5_-47; replicates of nanodisc formation with AASTY_8.9_-45; absorption spectrum of select oCPC samples and determination of a dissociation constant between Ca^2+^ and AASTY_7.4_-52; determination of wavelength to be used for analysis of copolymers by absorption; additional size exclusion chromatograms of pure copolymer and nanodiscs with CaCl_2_ or MgCl_2_; analysis of nanodisc size and peak shape for the different copolymers; size exclusion chromatograms of dialyzed nanodiscs of AASTY_8.9_-45 or AASTY_7.4_-52 diluted into buffers with CaCl_2_ or MgCl_2_; size exclusion chromatograms of different lipid concentrations solubilized by AASTY_8.9_-45; absorption spectra for quantification of free copolymer fractions in nanodisc samples of AASTY_12.5_-47(-ttc) and AASTY_7.4_-52(-ttc); free copolymer fraction as a function of lipid concentration for various nanodisc preparations; and absorption spectra of copolymer and nanodisc samples from SEC.

## Acknowledgement

The authors thank the Novo Nordisk Foundation (NNF20OC0060692) and the Carlsberg Foundation (CF20-0533) for support to equipment and infrastructure. A.A.A.A. was funded by grants NNF18OC0030896 from the Novo Nordisk Foundation and the Stanford Bio-X Program and 0171-00081B from Independent Research Fund Denmark. H.E.A. was funded by grant R265-2017-4015 from the Lundbeck Foundation. Subsets of the figures were created with BioRender.com. The authors thank Casper De Lichtenberg, Matilde Knapkøien Nor-dentoft and Sigrid Egevang Jensen for discussion of data and proofreading the manuscript.

## Graphical TOC Entry

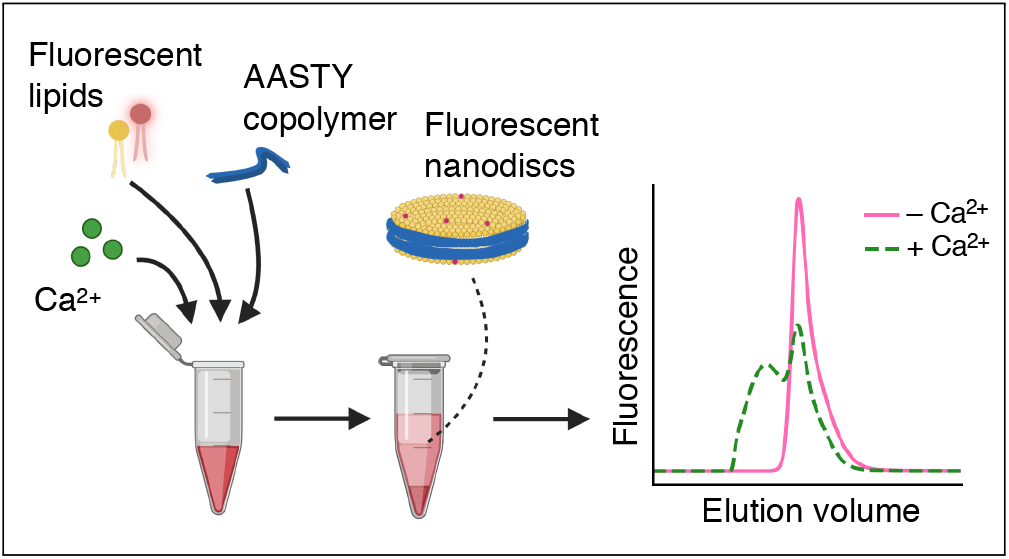

