## Supplemental_Information for "Characterization of divalent cation interactions with AASTY native nanodiscs"

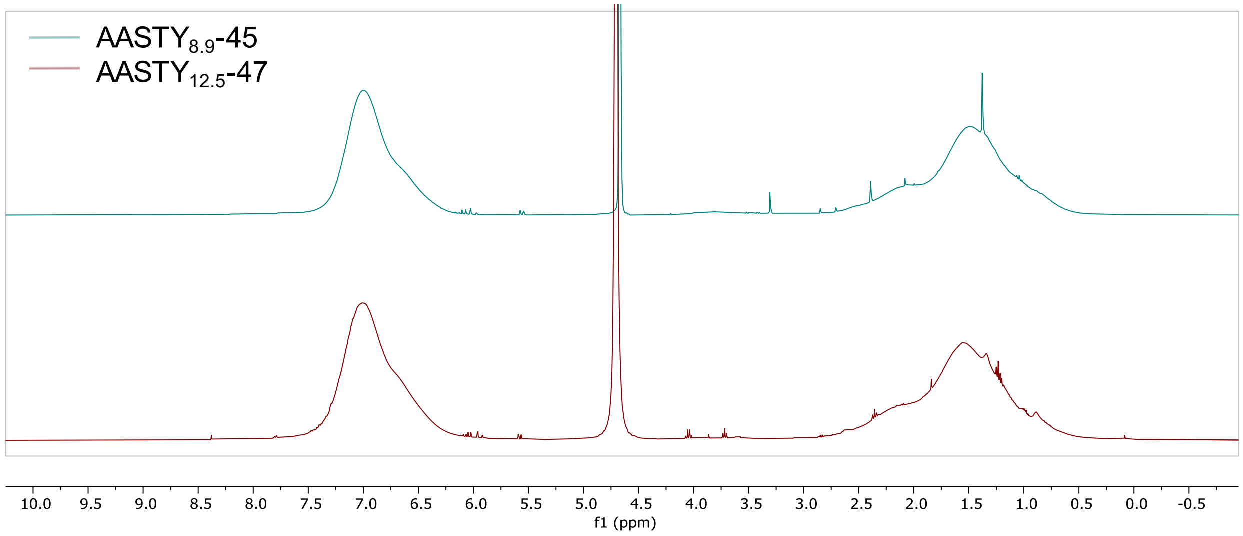


Figure S1: NMR spectrum of newly synthesized AASTY$_{12.5}$-47 compared with that of AASTY$_{8.9}$-45.[1] Phenyl protons are seen at 7.5-6.2 ppm, and backbonse protons at 3.0-0.5 ppm.

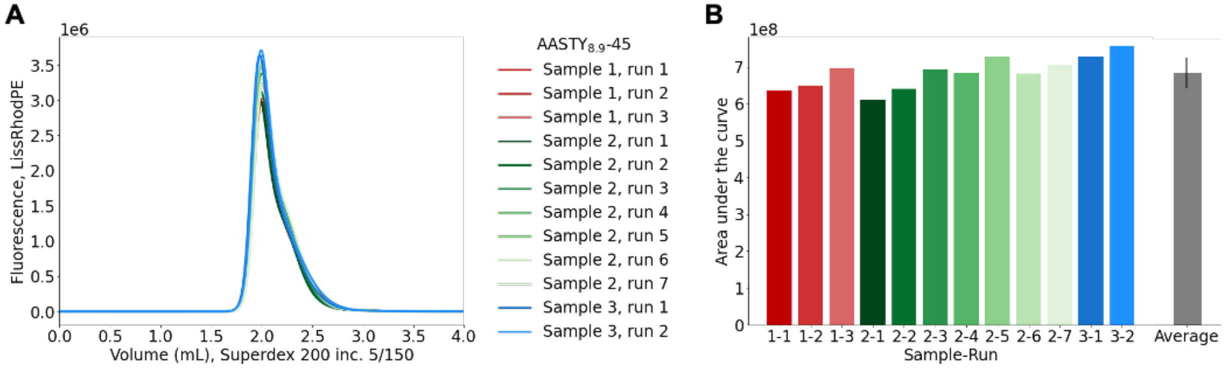


Figure S2: **A)** Overlay of FSEC traces of nanodiscs prepared from 1 mM 98 % POPC and 2 % LissRhodPE solubilized with 1 % AASTY$_{8.9}$-45 in 20 mM Hepes/NaOH, pH 7.4, 100 mM KCl without any $CaCl_2$ or $MgCl_2$. FSEC was run in the same buffer. Three individual solubilizations were each run on the column multiple times. **B)** Area under the curve of for the traces from A) as well as their average with the standard deviation.

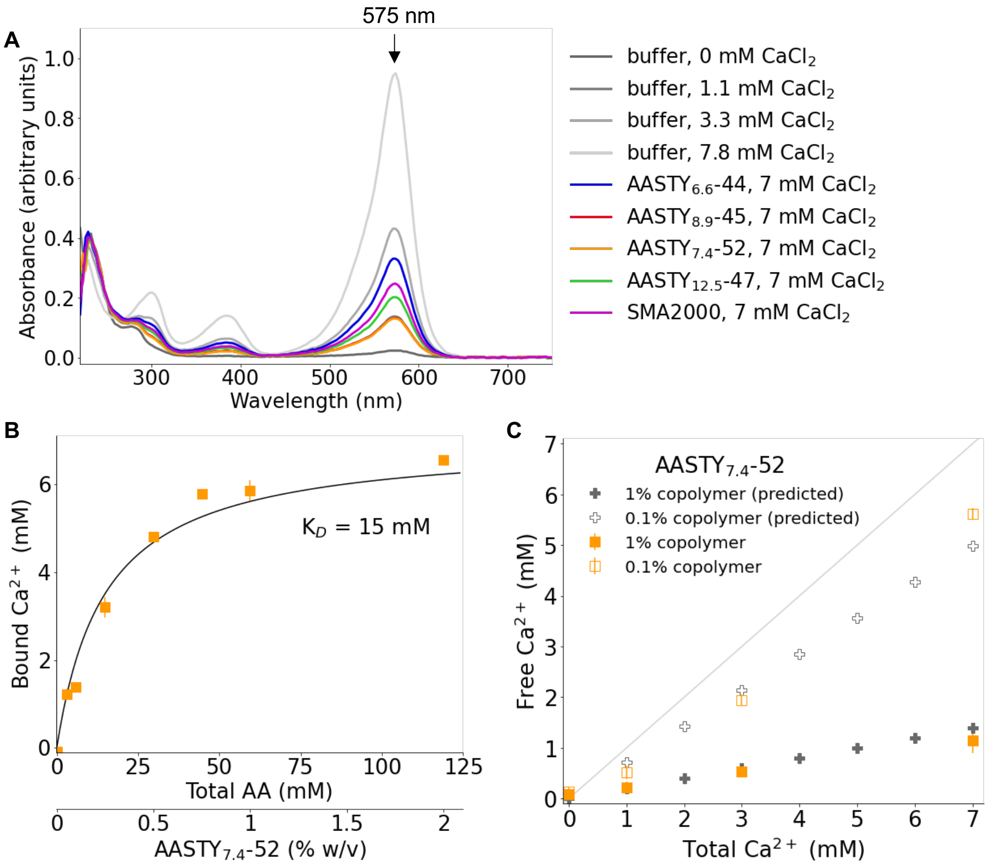


Figure S3: **A)** Spectra of select samples from the oCPC based free $Ca^{2+}$ assay in Figure 2B indicating the maximum absorbance at 575 nm. There is no difference between the buffer samples (greys) and those with copolymer at short wavelengths (<300 nm) where copolymers absorb strongly, showing that the copolymer has succesfully been removed in the centrifugal concentrator. **B)** Measurement of free $Ca^{2+}$ concentrations in the presence of 7 mM $CaCl_2$ and varying amounts of $AASTY_{7.4}$-52 using the oCPC based assay, shown as the amount of bound $Ca^{2+}$. To estimate the dissociation constant, $K_D$, the binding curve was fit with the approximation that all AA groups could be treated as free. Each measurement was performed in duplicate, apart from 1 % and 0.1 % copolymer, which were performed in triplicate. **C)** Measurement of free $Ca^{2+}$ concentrations in the presence of 1 % and 0.1 % $AASTY_{7.4}$-52 compared with values calculated using the $K_D$ determined in B). Each condition was assayed in triplicates. The 1 % data is also shown in Figure 2B.

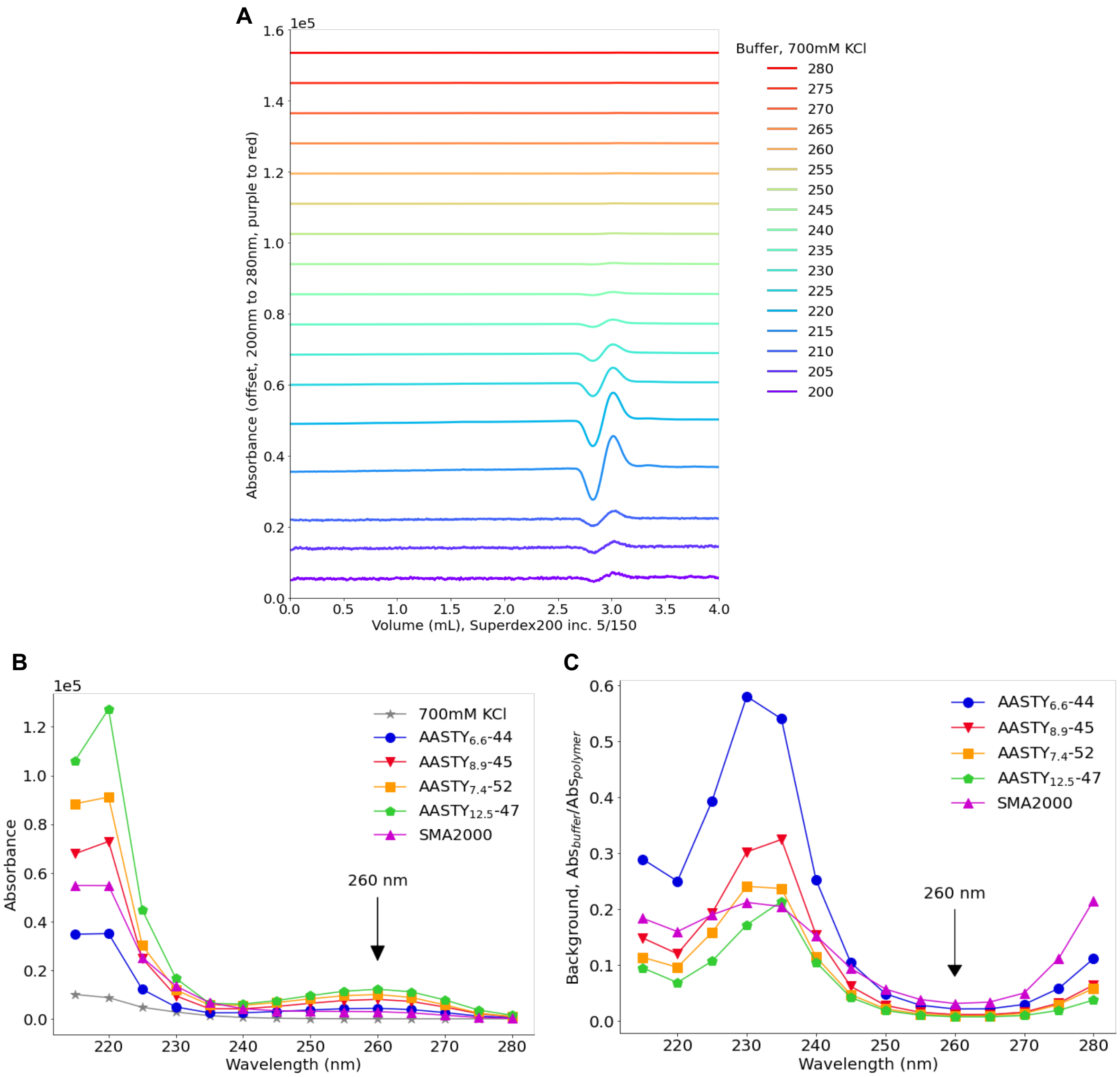


Figure S4: **A)** Spectrum of loading 2 μL buffer with 700 mM KCl, in running buffer with only 100 mM KCl. Analogous spectra for the five analyzed copolymers are found in Supp. Figures S14-S18. **B)** Maximum absorbance at different wavelengths for the buffer from A) and for loading 2 μL 0.1 % pure copolymer. **C)** The potential "background" signal from the 700 mM KCl buffer estimated as the maximum absorbance of the buffer divided by the maximum absorbance of the copolymer calculated at specified wavelengths. 260 nm was chosen for analysis of copolymers, as the effect of buffer should be minimal, while it is also a local maximum for AASTY copolymers (see B).

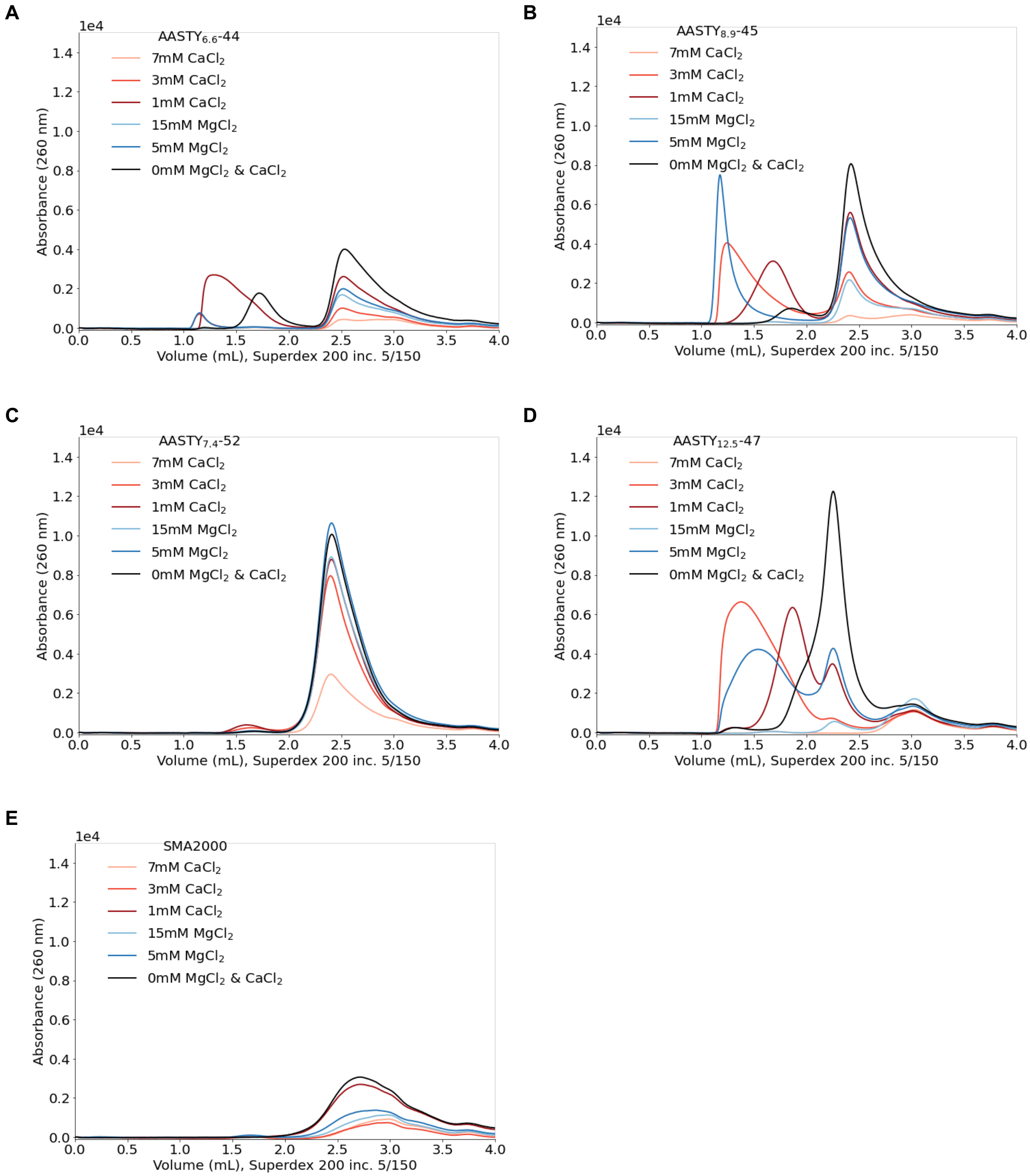


Figure S5: Size exclusion chromatograms of 0.1 % copolymer in the presence of different concentrations of either $CaCl_2$ or $MgCl_2$. The running buffer in all instances was 20 mM Hepes/NaOH, pH 7.4, 100 mM KCl without any $CaCl_2$ or $MgCl_2$. The area under the curve for these traces is quantified in Figure 2E. **A)** AASTY$_{6.6}$-44, **B)** AASTY$_{8.9}$-45, **C)** AASTY$_{7.4}$-52, **D)** AASTY$_{12.5}$-47, and **E)** SMA2000.

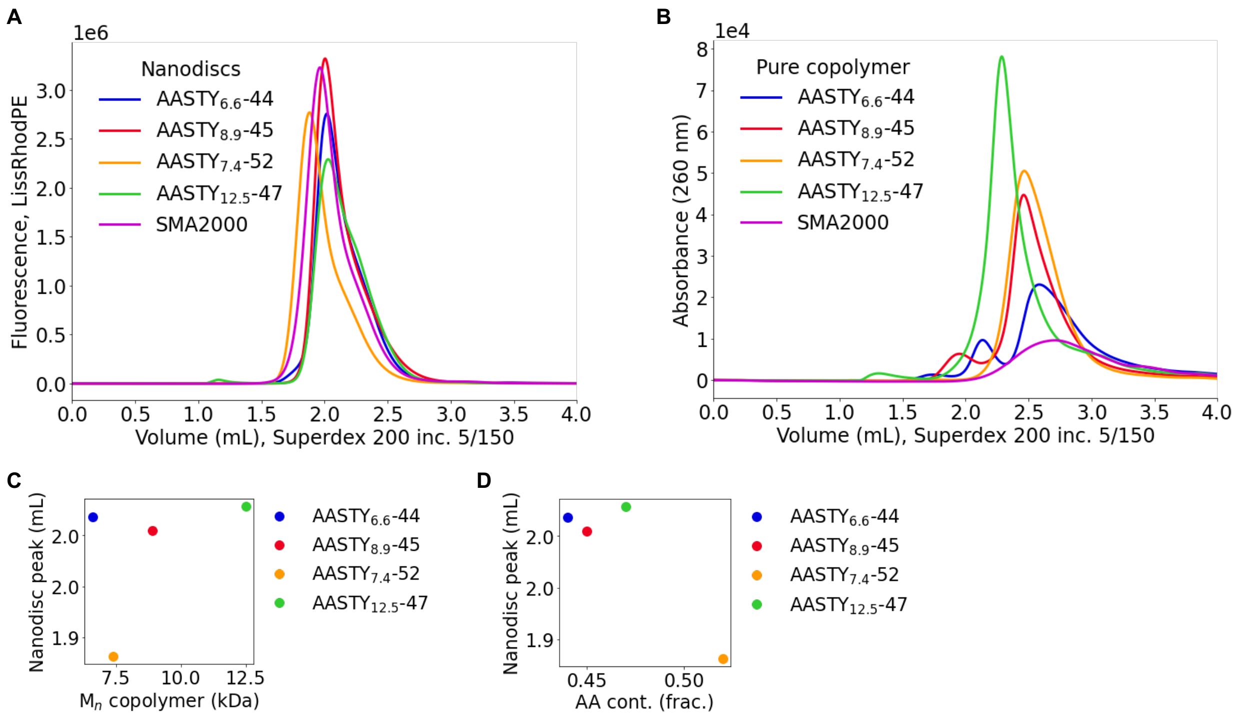


E

| Copolymer | Position of peak (mL) | Height of peak | Area under the curve | Full width at half max (mL) | Full width at quarter max (mL) | FWQM-FWHM (mL) |
|---|---|---|---|---|---|---|
| $AASTY_{6.6}$-44 | 2.02 | 2.86e6 | 6.01e8 | 0.308 | 0.517 | 0.218 |
| $AASTY_{8.9}$-45 | 2.01 | 3.32e6 | 6.82e8 | 0.258 | 0.485 | 0.227 |
| $AASTY_{7.4}$-52 | 1.88 | 2.77e6 | 5.87e8 | 0.272 | 0.510 | 0.238 |
| $AASTY_{12.5}$-47 | 2.03 | 2.29e6 | 5.79e8 | 0.395 | 0.583 | 0.188 |
| SMA2000 | 1.96 | 3.23e6 | 7.13e8 | 0.283 | 0.525 | 0.241 |

Figure S6: The size of nanodiscs does not correlate with the molecular weight of the copolymer used. **A)** Nanodiscs produced by solubilizing 1 mM 98 % POPC and 2 % LissRhodPE with 1 % of indicated copolymer (0 mM $MgCl_2$ and $CaCl_2$ traces from Supp. Fig. S7 overlayed). **B)** Comparison of SEC traces from loading 1 % copolymer (Supp. Fig. S14-S18A). **C)** Number average molecular weight ($M_n$ vs the position of the nanodisc peak for the AASTY nanodiscs. Lower elution volume translates to larger nanodiscs. **D)** Acrylic acid (AA) content vs the position of the nanodisc peak for the AASTY nanodiscs. **E)** Table with quantification of the nanodisc traces from A). The difference between the full width at half max (FWHM) and the full width at quarter max (FWQM) can indicate if the peak has a shoulder.

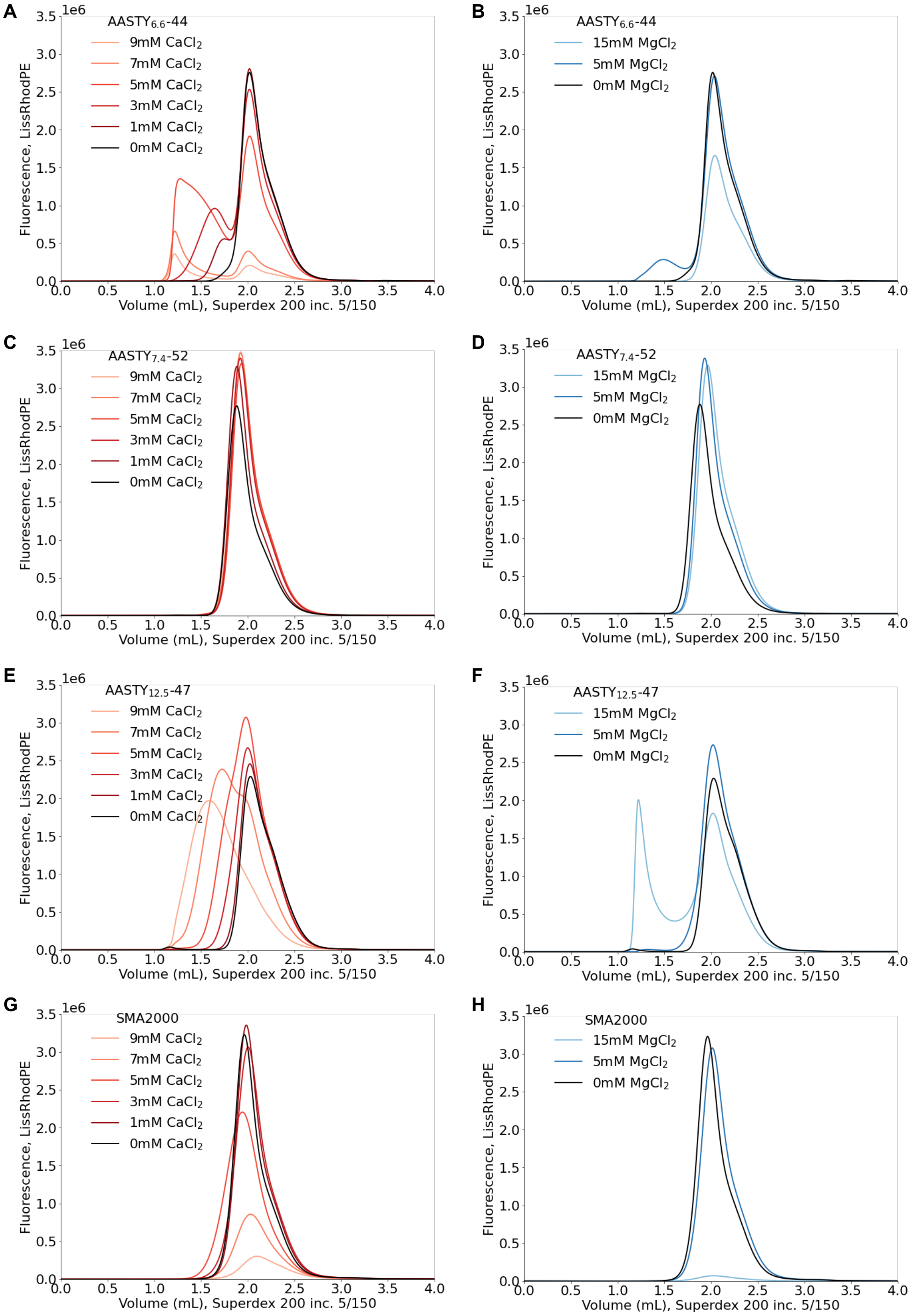


Figure S7: Solubilization of small unilamellar vesicles (SUVs) of 98 % POPC and 2 % LissRhodPE by copolymers in the presence of different concentrations of either $CaCl_2$ or $MgCl_2$. The running buffer in all instances was 20 mM Hepes/NaOH, pH 7.4, 100 mM KCl without any $CaCl_2$ or $MgCl_2$. Nanodiscs are visualized by the fluorescence of LissRhodPE. The area under the curve is quantified in Figure 3C, and the SEC profiles for $AASTY_{8.9}$-45 are shown in Figure 3A,B. **A,B)** $AASTY_{6.6}$-44, **C,D)** $AASTY_{7.4}$-52, **E,F)** $AASTY_{12.5}$-47, and **G,H)** SMA2000.

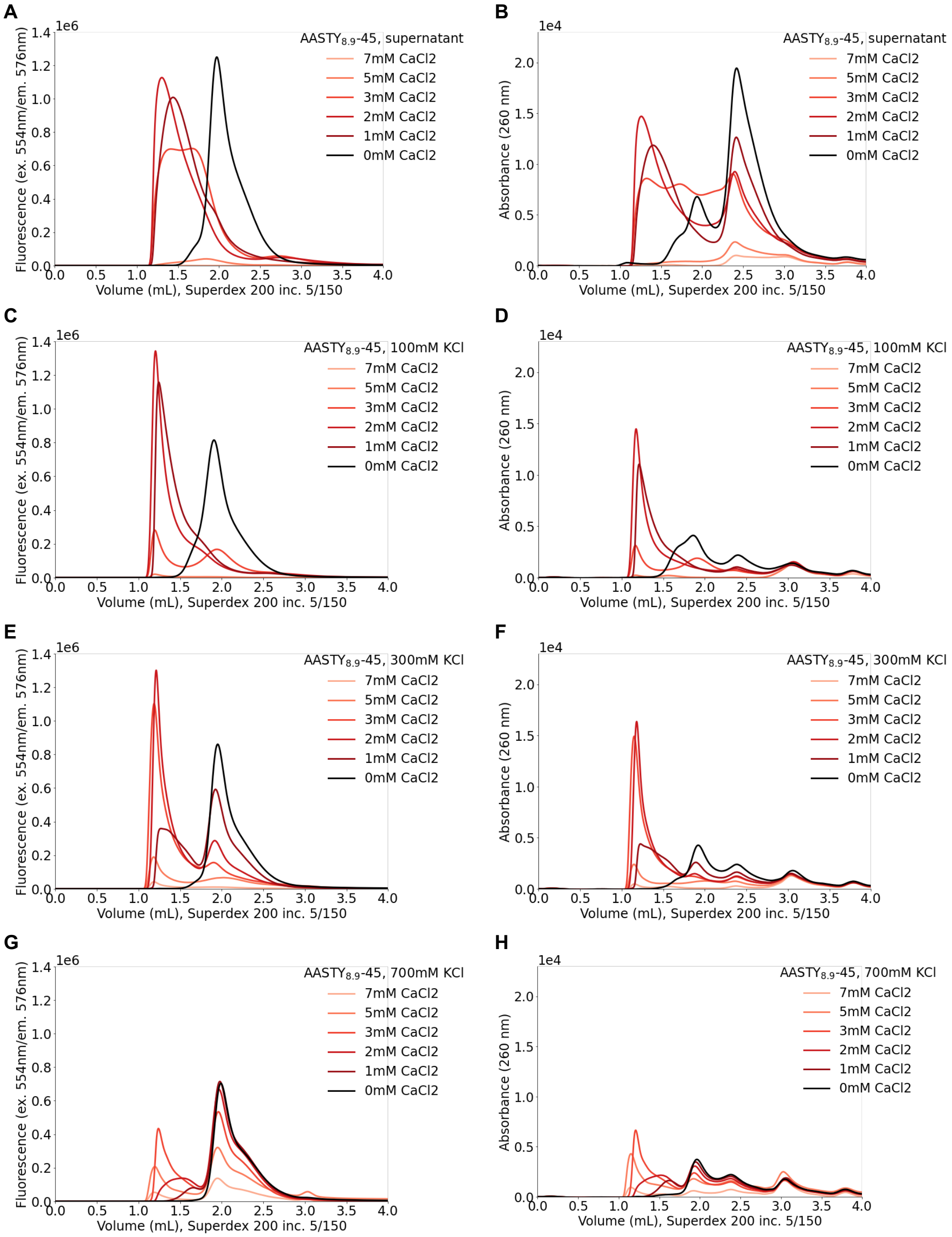


Figure S8: AASTY$_{8.9}$-45 nanodiscs were prepared from SUVs of 98 % POPC and 2 % LissRhodPE and diluted 10-fold into specified buffers. The running buffer in all instances was 20 mM Hepes/NaOH, pH 7.4, 100 mM KCl without any $CaCl_2$. **A,C,E,G)** Nanodiscs visualized by LissRhodPE fluorescence. **B,D,F,H)** copolymer visualized at absorbance of 260 nm. The areas under the curve are quantified in Figures 4B and 6C. **A,B)** Samples that were not dialyzed diluted in 100 mM KCl with specified $CaCl_2$. **C-H)** Samples dialyzed to remove free copolymer diluted in buffer with specified KCl and $CaCl_2$ concentrations.

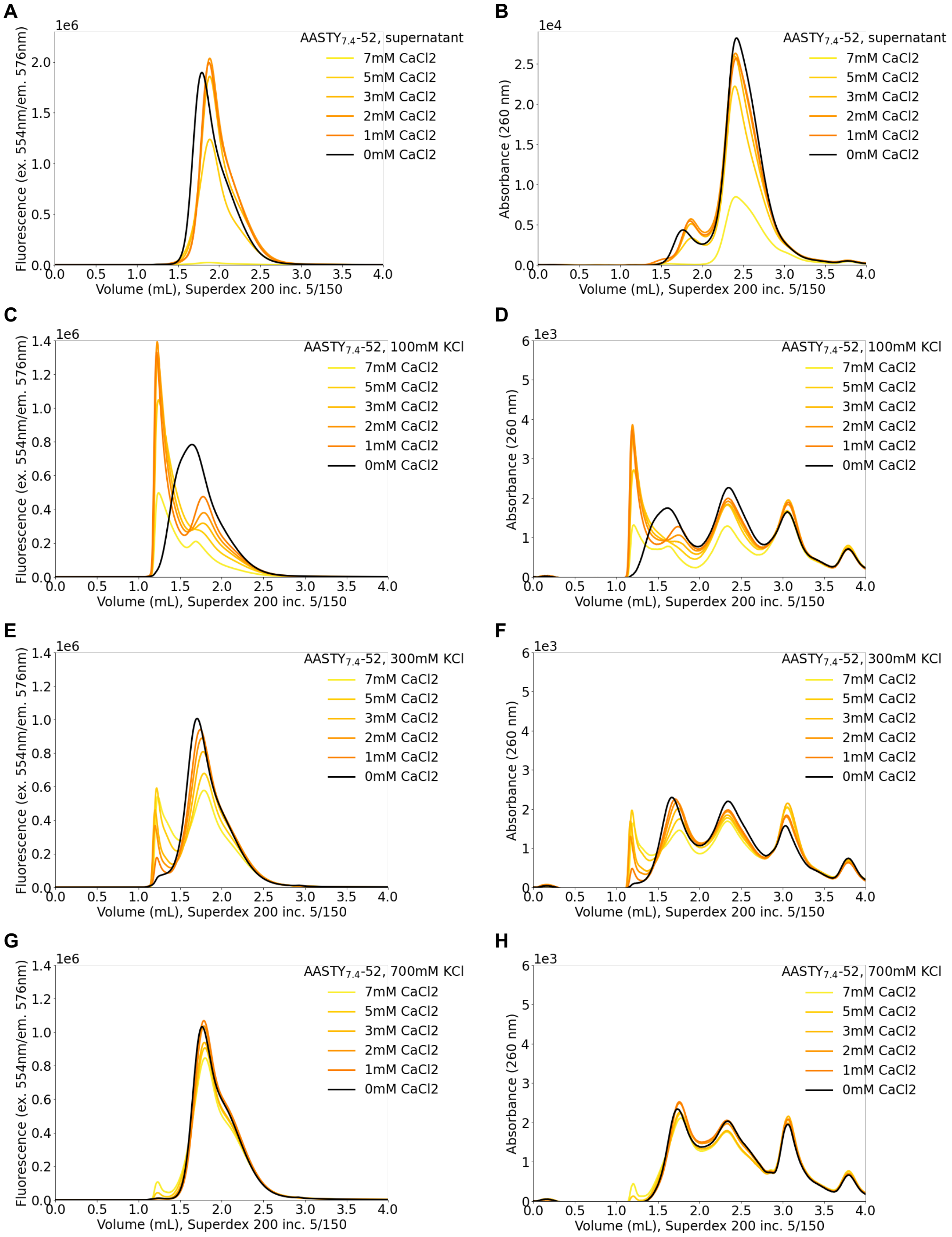


Figure S9: AASTY$_{7.4}$-52 nanodiscs were prepared from SUVs of 98 % POPC and 2 % LissRhodPE and diluted 10-fold into specified buffers. The running buffer in all instances was 20 mM Hepes/NaOH, pH 7.4, 100 mM KCl without any $CaCl_2$. **A,C,E,G)** Nanodiscs visualized by LissRhodPE fluorescence. **B,D,F,H)** copolymer visualized at absorbance of 260 nm. The areas under the curve are quantified in Figures 4C and 6E. **A,B)** Samples that were not dialyzed diluted in 100 mM KCl with specified $CaCl_2$. **C-H)** Samples dialyzed to remove free copolymer diluted in buffers with specified KCl and $CaCl_2$ concentrations.

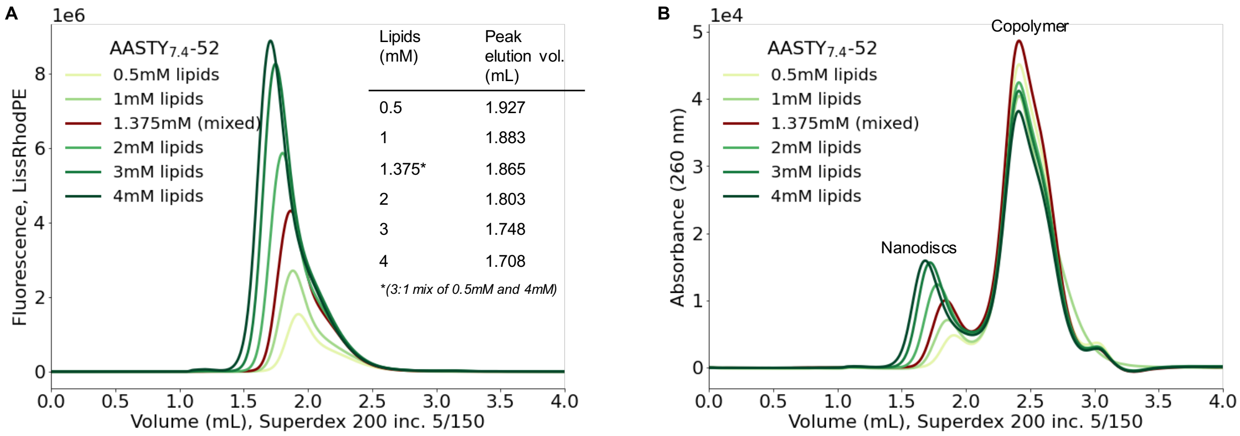


Figure S10: **A)** FSEC traces of nanodiscs generated by solubilizing different total lipid concentrations (98 % POPC and 2 % LissRhodPE) with 1 % AASTY$_{7.4}$-52. The table insert indicates the elution volume of the peak for different samples. The sample with 1.375 mM lipids (brown trace) was produced by mixing the 0.5 mM and 4 mM samples. **B)** A$_{260}$ for the samples in A). Quantification of areas under the curve and comparison with AASTY$_{8.9}$-45 can be found in Figure 7.

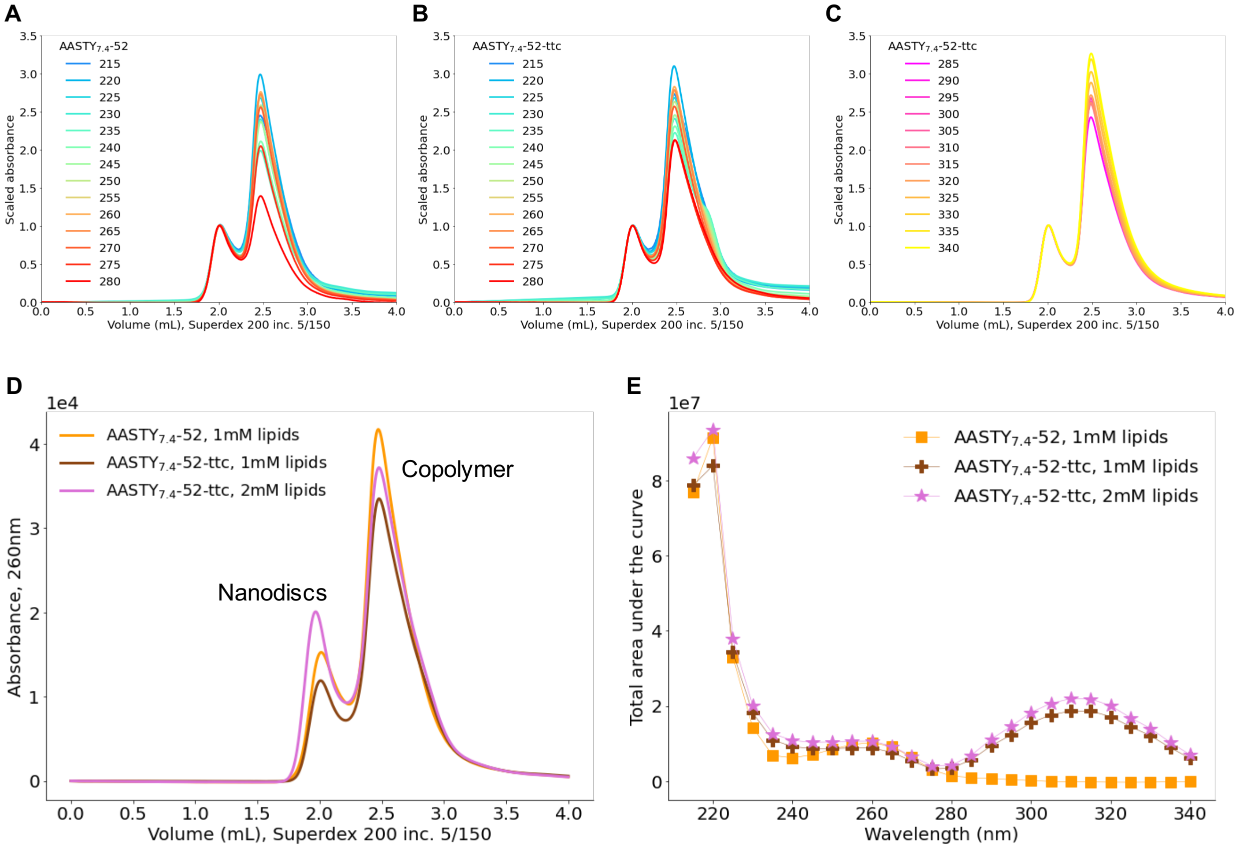


Figure S11: **A)** Absorbance spectrum of the AASTY$_{7.4}$-52 trace from C) at 215-280 nm, normalized by the nanodisc peak to illustrate the difference in the spectral properties of nanodiscs and free copolymer. **B)** Like A) for AASTY$_{7.4}$-52-ttc with 1 mM lipids. **C)** Wavelengths (285-340 nm) dominated by the ttc group for AASTY$_{7.4}$-52-ttc with 1 mM lipids, normalized as D). **D)** SEC traces of solubilizing 1 or 2 mM SUVs of 98% POPC and 2% LissRhodPE with 1% AASTY$_{7.4}$-52 with or without the ttc tag. **E)** Maximum absorbance at different wavelengths for the traces from D). The ttc group has a maximum absorbance at 310 nm.

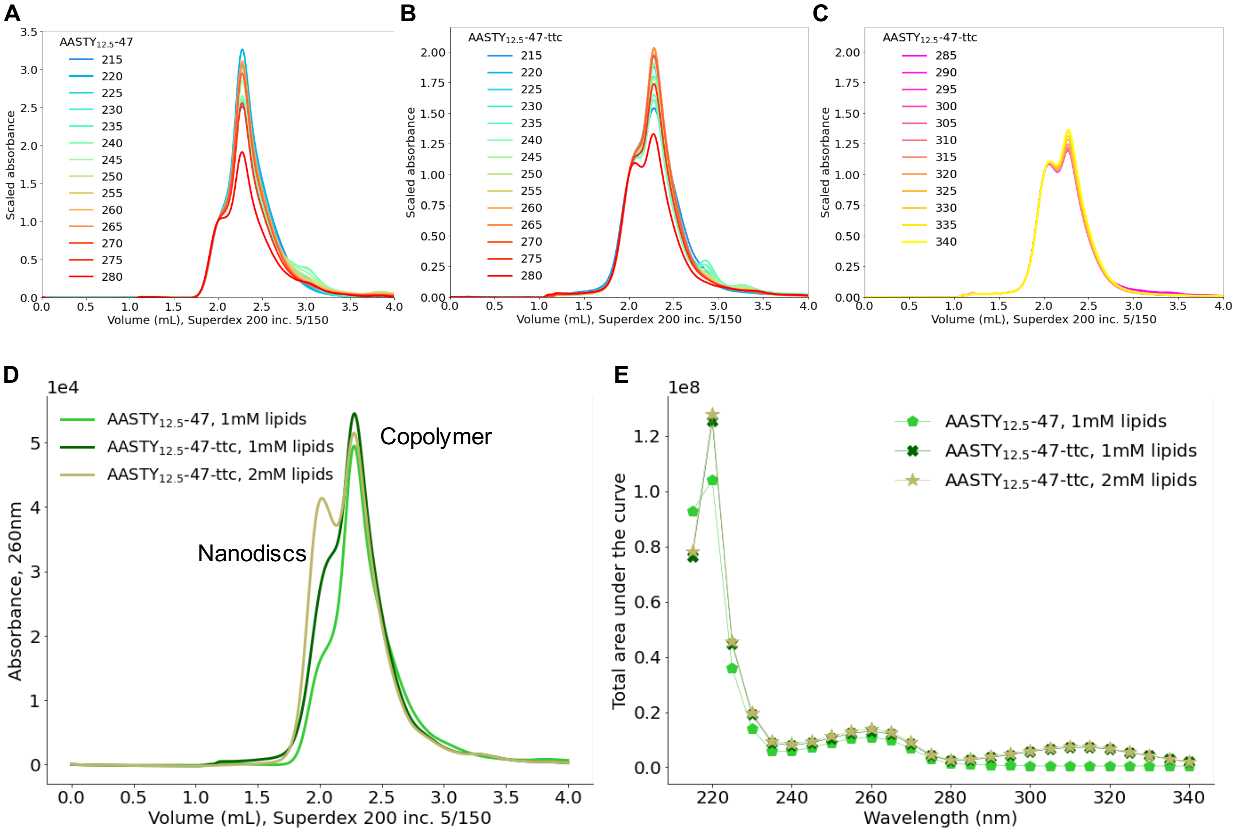


Figure S12: **A)** Absorbance spectrum of the AASTY$_{12.5}$-47 trace from C) at 215-280 nm, normalized by the nanodisc peak to illustrate the difference in the spectral properties of nanodiscs and free copolymer. **B)** Like A) for AASTY$_{12.5}$-47-ttc with 1 mM lipids. **C)** Wavelengths (285-340 nm) dominated by the ttc group for AASTY$_{12.5}$-47-ttc with 1 mM lipids, normalized as D). **D)** SEC traces of solubilizing 1 or 2 mM SUVs of 98% POPC and 2% LissRhodPE with 1% AASTY$_{12.5}$-47 with or without the ttc tag. **E)** Maximum absorbance at different wavelengths for the traces from D). The ttc group has a maximum absorbance at 310 nm.

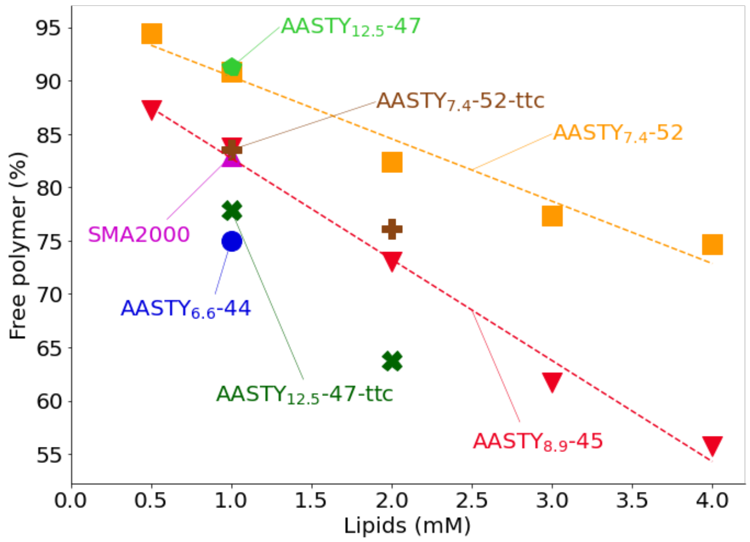


Figure S13: Percentage of free copolymer estimated at 260 nm for copolymers without the ttc group, and 310 nm for AASTY$_{7.4}$-52-ttc and AASTY$_{12.5}$-47-ttc as a function of the lipid concentration present during solubilization. For AASTY$_{8.9}$-45 and AASTY$_{7.4}$-52(-ttc) four Gaussian functions were used for fitting, while three were used for the rest.

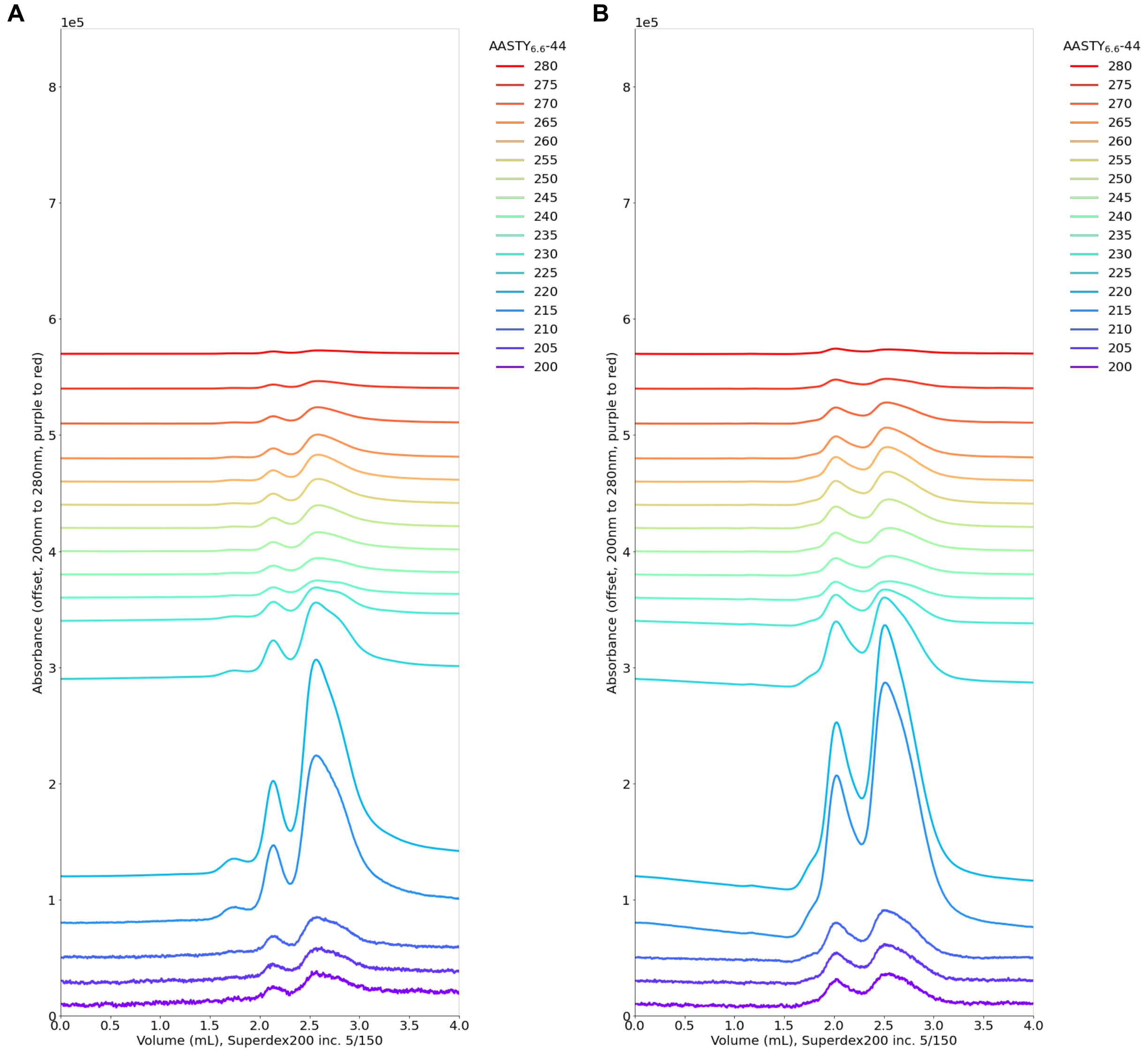


Figure S14: Absorption spetrum of AASTY$_{6.6}$-44 **A)** 1 % copolymer and **B)** nanodisc prepared from 98 % POPC and 2 % LissRhodPE. The colors indicate absorbance at different wavelengths. For both 2 µL were loaded.

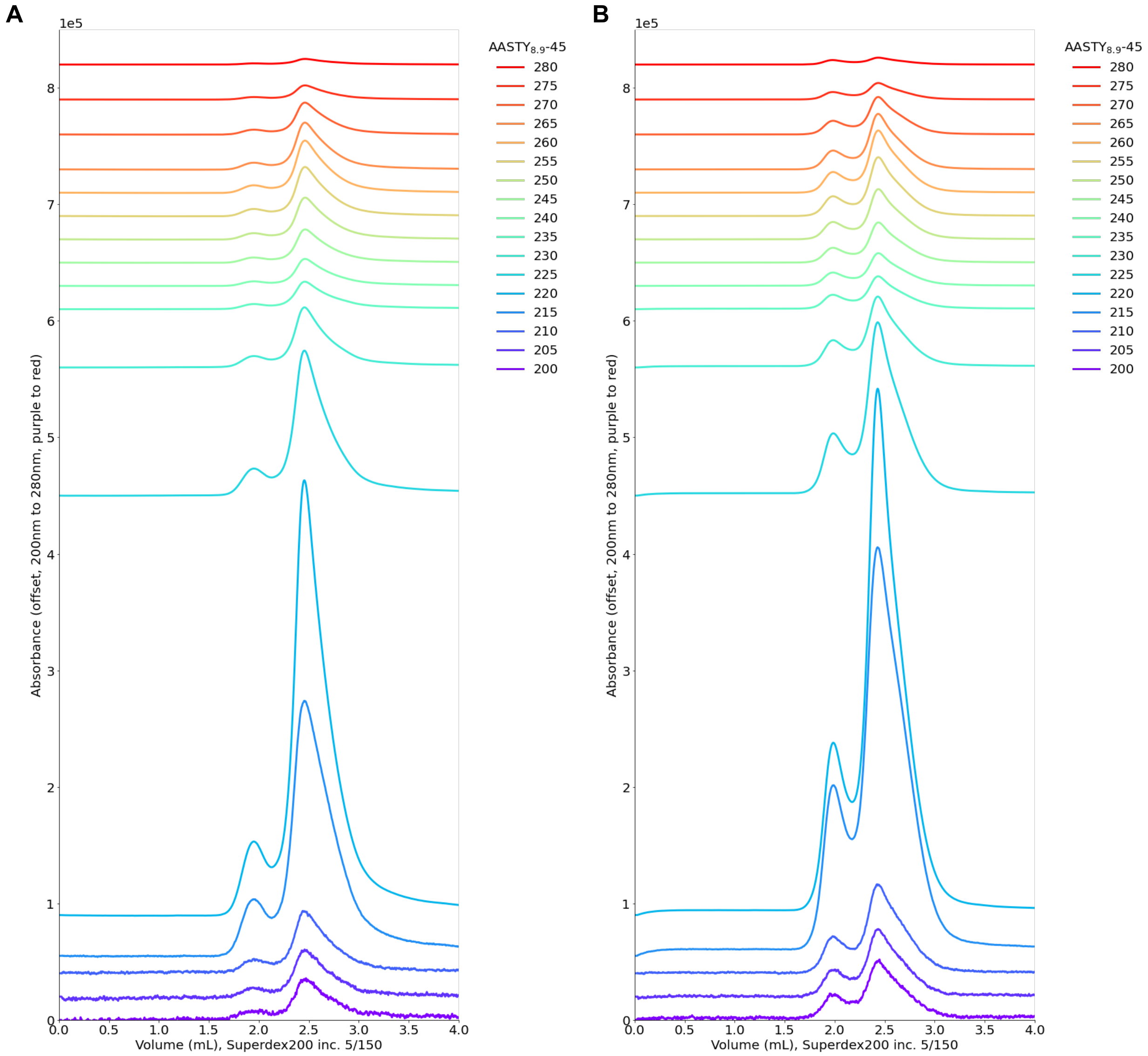


Figure S15: Absorption spetrum of AASTY$_{8.9}$-45 **A)** 1 % copolymer and **B)** nanodisc prepared from 98 % POPC and 2 % LissRhodPE. The colors indicate absorbance at different wavelengths. For both 2 µL were loaded.

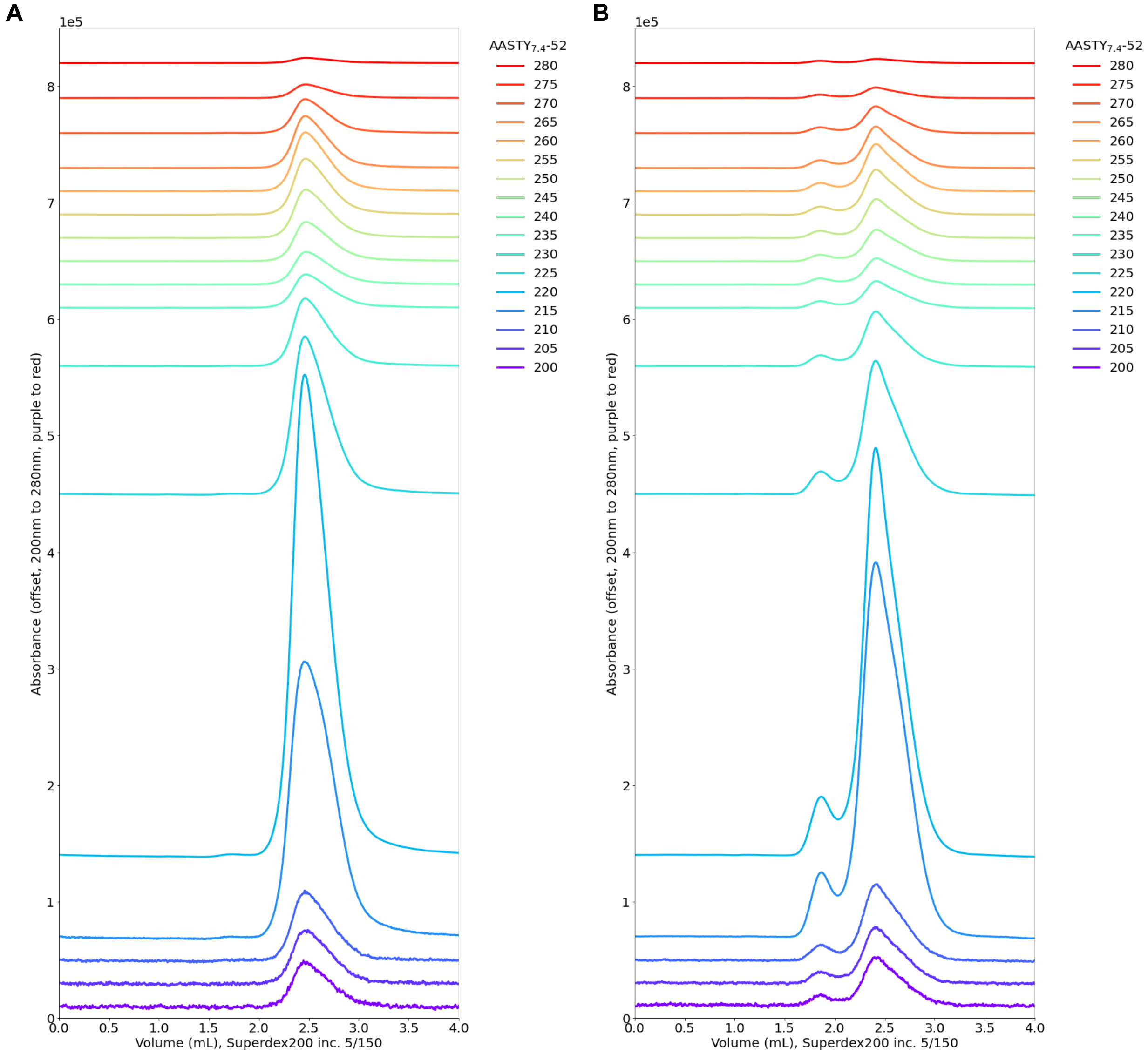


Figure S16: Absorption spetrum of AASTY$_{7.4}$-52 **A)** 1 % copolymer and **B)** nanodisc prepared from 98 % POPC and 2 % LissRhodPE. The colors indicate absorbance at different wavelengths. For both 2 µL were loaded.

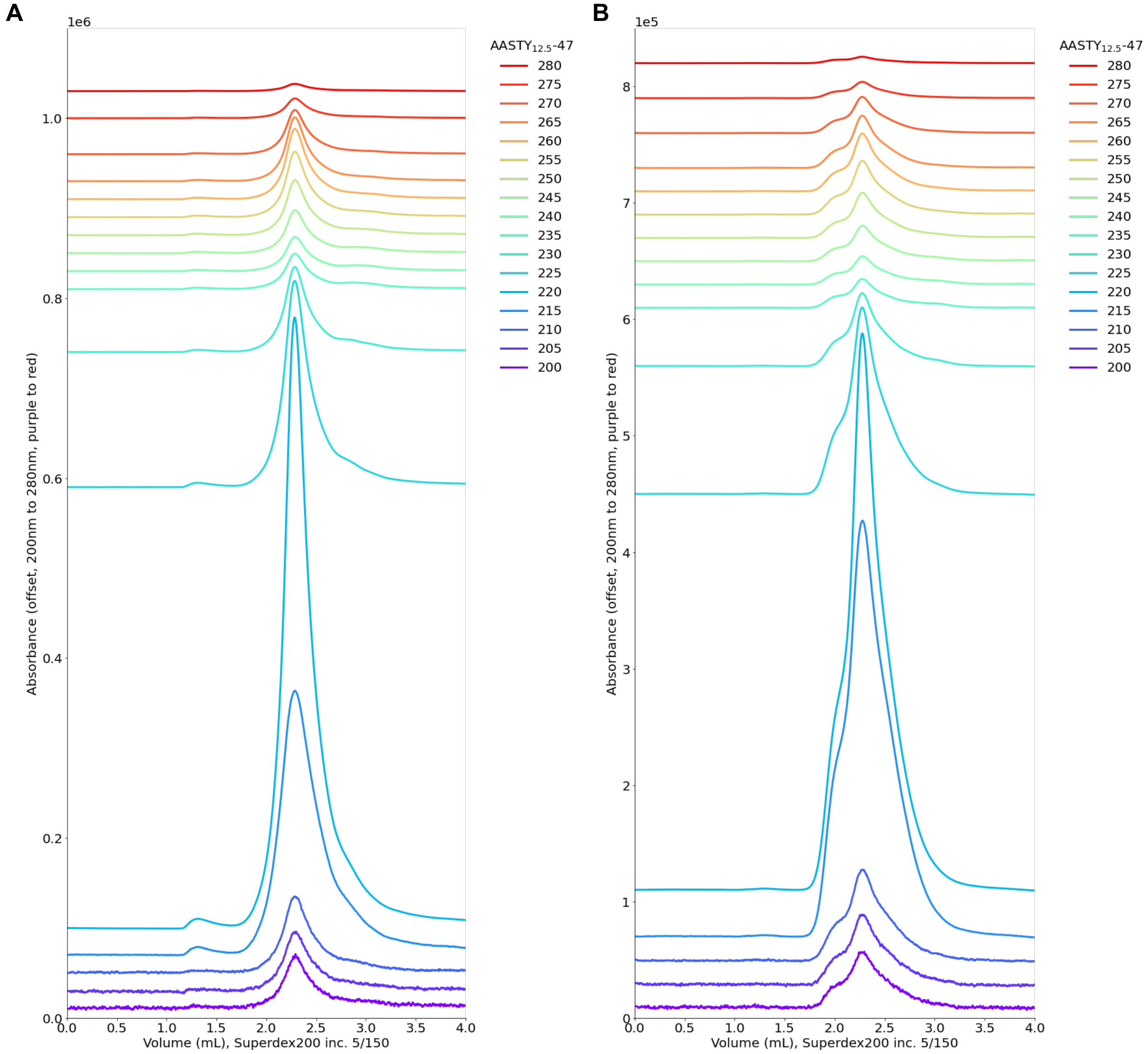


Figure S17: Absorption spetrum of AASTY$_{12.5}$-47 **A)** 1 % copolymer and **B)** nanodisc prepared from 98 % POPC and 2 % LissRhodPE. The colors indicate absorbance at different wavelengths. For both 2 µL were loaded.

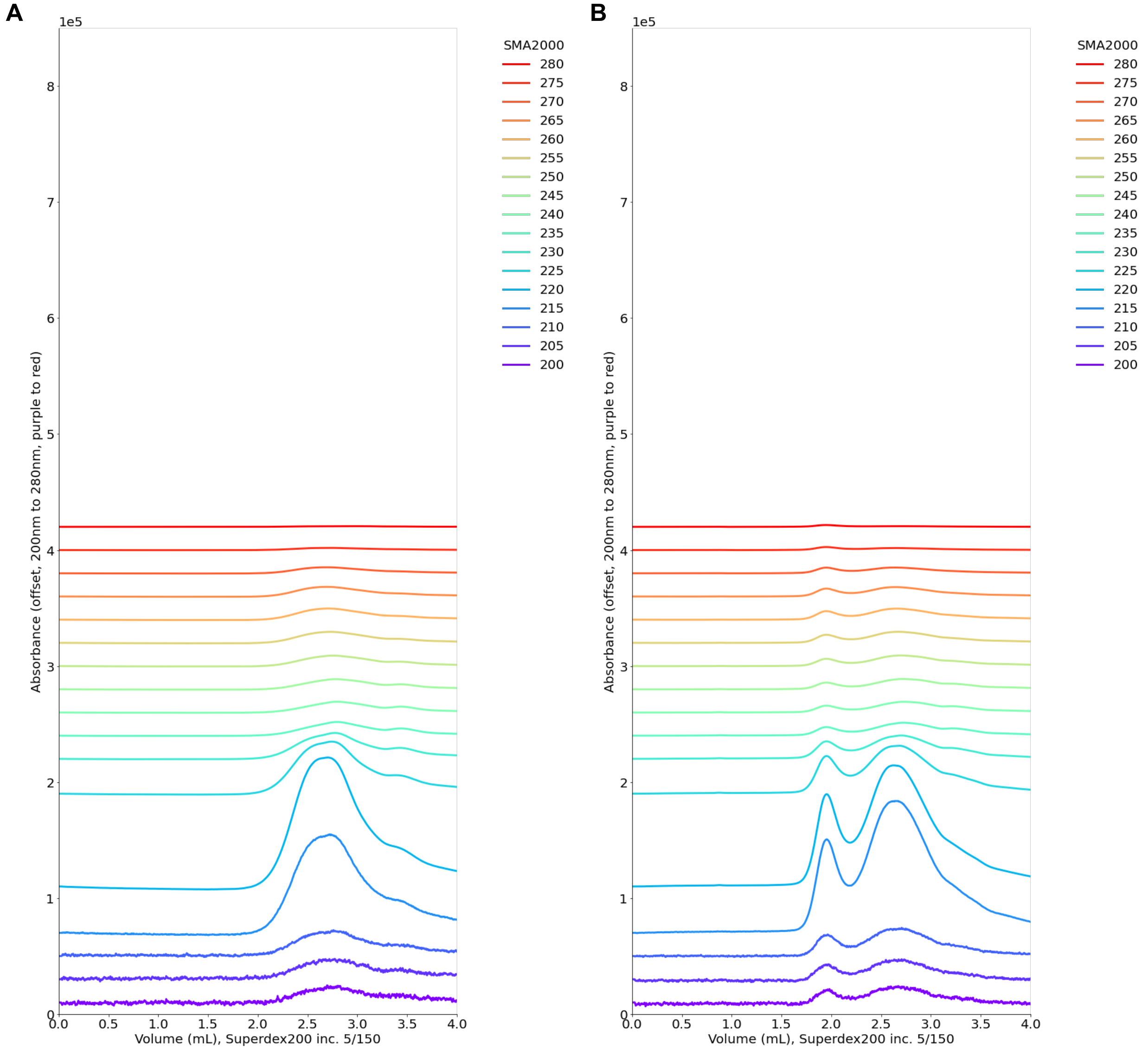


Figure S18: Absorption spetrum of SMA2000 **A)** 1 % copolymer and **B)** nanodisc prepared from 98 % POPC and 2 % LissRhodPE. The colors indicate absorbance at different wavelengths. For both 2 µL were loaded.
